# Ecological load and balancing selection in circumboreal barnacles

**DOI:** 10.1101/2020.07.18.209569

**Authors:** Joaquin C. B. Nunez, Stephen Rong, Alejandro Damian-Serrano, John T. Burley, Rebecca G. Elyanow, David A. Ferranti, Kimberly B. Neil, Henrik Glenner, Magnus Alm Rosenblad, Anders Blomberg, Kerstin Johannesson, David M. Rand

## Abstract

Acorn barnacle adults experience environmental heterogeneity at various spatial scales of their circumboreal habitat, raising the question of how adaptation to high environmental variability is maintained in the face of strong juvenile dispersal and mortality. Here we show that 4% of genes in the barnacle genome experience balancing selection across the entire range of the species. Many of these genes harbor mutations maintained across 2 million years of evolution between the Pacific and Atlantic oceans. These genes are involved in ion regulation, pain reception, and heat tolerance, functions which are essential in highly variable ecosystems. The data also reveal complex population structure within and between basins, driven by the trans-Arctic interchange and the last glaciation. Divergence between Atlantic and Pacific populations is high, foreshadowing the onset of allopatric speciation, and suggesting that balancing selection is strong enough to maintain functional variation for millions of years in the face of complex demography.

## Introduction

The relationship between genetic variation and adaptation to heterogeneous environments remains a central conundrum in evolutionary biology (Botero, et al. 2015). Classical models of molecular evolution predict that populations should be locally adapted to maximize fitness (Williams 1966). However, species inhabiting highly heterogeneous environments violate this expectation: if gene flow is high in relation to the scale of environmental heterogeneity, species will harbor variation that is beneficial in one condition but deleterious in another (Gillespie 1973), and the resulting ecological load (i.e., the fitness difference between the best and the average genotype across the range of environments where offspring may settle) will prevent local adaptation. Conversely, if gene flow is low, favored alleles will become locally fixed and species should display low levels of genetic variation. Paradoxically, many natural populations living in variable environments possess high dispersal capabilities, and harbor more variation than expected under classical models (Metz and Palumbi 1996; Mackay, et al. 2012; Messer and Petrov 2013; Bergland, et al. 2014). This disconnect between nature and theory has motivated the hypothesis that balancing selection, a process where selection favors multiple beneficial alleles at a given locus, is at play to maintain adaptations in these habitats (Levene 1953; Hedrick 2006).

The northern acorn barnacle (*Semibalanus balanoides*) is a model system to study adaptations to ecological variability. This barnacle is a self-incompatible, simultaneous hermaphrodite which outcross only with adjacent individuals. Adult barnacles are fully sessile and occupy broad swaths of intertidal shores in both the North Pacific and North Atlantic oceans. These habitats experience high levels of cyclical and stochastic ecological heterogeneity which impose strong selection at multiple spatial scales: microhabitats (intertidal shores), mesohabitats (bays and estuaries) and macrohabitats (continental seaboards) (Schmidt, et al. 2008; Nunez, et al. 2020). Barnacle larvae, on the other hand, engage in extensive pelagic dispersal by ocean currents, and may settle in habitats completely different from those of their parents (Flowerdew 1983). This contrast between strong adult selection and high juvenile dispersal prevents local adaptation. In addition, *S. balanoides* has a complex demography. It originated in the Pacific, and colonized the Atlantic during the many waves of the trans-Arctic interchange (1-3 mya) (Vermeij 1991). Like most circumboreal species, it was subjected to drastic range shifts due to the Pleistocene glacial cycles (Wares and Cunningham 2001; Flight, et al. 2012), and more recently due to anthropogenic climate change (Jones, et al. 2012). As such, S. *balanoides* is a premier system to study how adaptive genetic variation is maintained over broad spatial and evolutionary scales, in the face of ecological load.

Three decades of work have shown that balancing selection, via marginal overdominance (a case where the harmonic mean fitness of heterozygous genotypes must be larger than that of either homozygote) (Levene 1953), maintains adaptive variation at the metabolic gene Mannose-6-phopate isomerase (***Mpi***) in barnacles across the entire North Atlantic basin(Schmidt and Rand 1999; Dufresne, et al. 2002; Rand, et al. 2002; Veliz, et al. 2004; Nunez, et al. 2020). These findings motivate two questions which are addressed in this paper. First, how pervasive are balanced polymorphisms in the barnacle genome? And, second, what genes are targets of balancing selection? To investigate functional polymorphism in *S. balanoides*, we quantified genomic variation in North Pacific and North Atlantic populations (**Figs. 1A-1C**). In the Pacific, we analyzed samples from British Columbia, Canada (**WCAN**) as well as a sample of the sister taxon *Semibalanus cariosus*. In the Atlantic, we analyzed samples from Maine (**ME**), Rhode Island (**RI**), Iceland (**ICE**), Norway (**NOR**), and the United Kingdom (**UK**). For all populations, we sequenced multiple libraries including: a single individual barnacle genome to ∼50X coverage, pools of 20-38 individuals per population (i.e., pool-seq (Schlotterer, et al. 2014)), as well as ∼600 bp amplicons from the mitochondrial (**mtDNA**) *COX I* gene (including previously published *COX I* data(Wares and Cunningham 2001)). We mapped these datasets to our newly assembled *S. balanoides* genome (**SI Appendix 1**) and characterized genetic diversity across all populations (**SI Appendix 2**). We first present our findings in the context of the barnacle’s phylogeography and demographic history. This is pivotal to understand the historical conditions which can contribute to ecological load. Then, we characterize the pervasiveness of balancing selection across the genome, as well as the age of balanced polymorphisms and their putative functional significance in highly heterogeneous environments.

**Figure 1.**
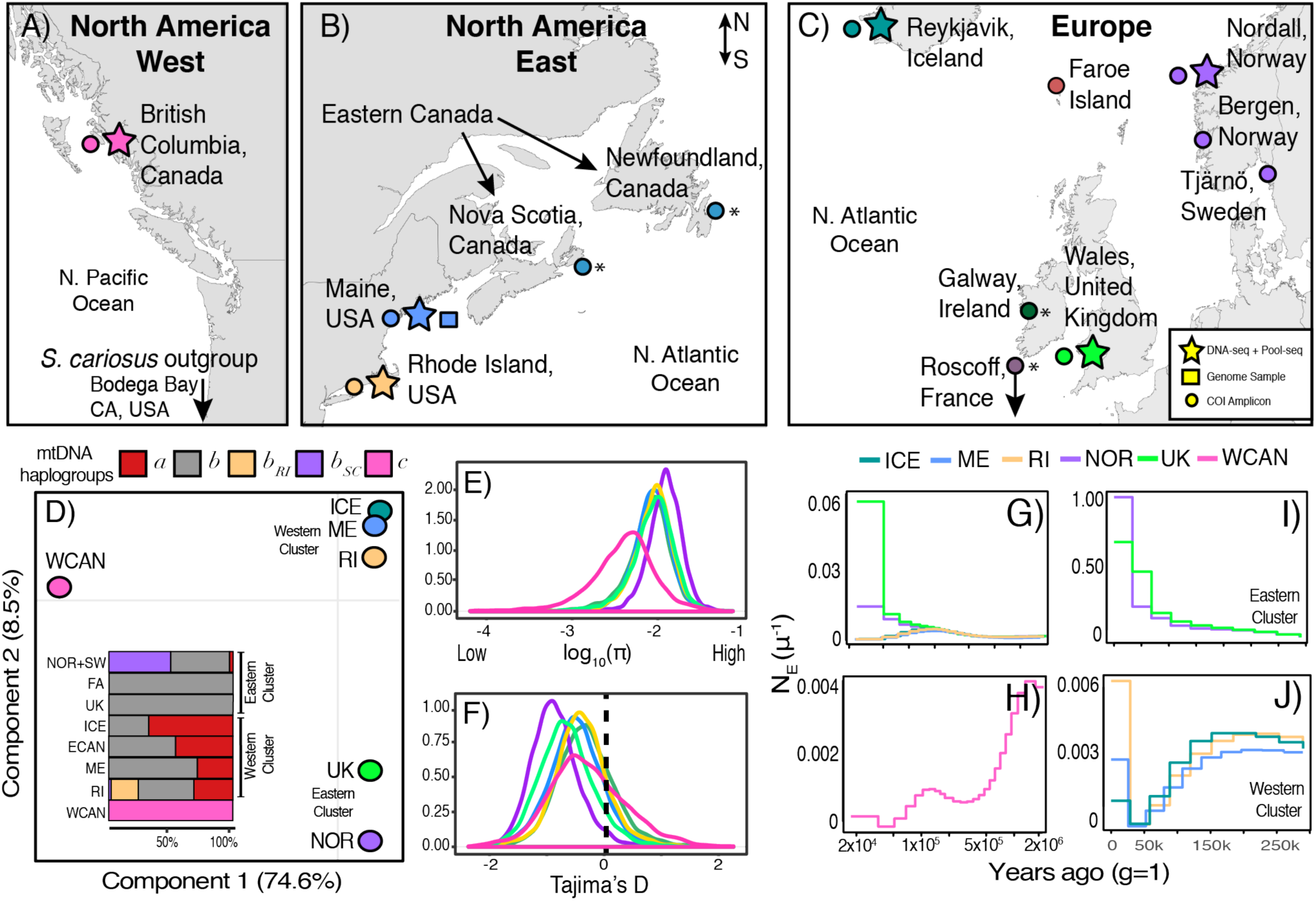
Genetic variation and phylogeography. **A**. Map of the North Pacific coast of North America with collection sites indicated. **B**. Collections in the Atlantic Eastern coast of North America. **C**. Collections in the Atlantic European coast. For A, B, and C, stars indicate sites where a single individual and a pool of multiple individuals were collected, the hexagon indicates the site from which the reference genome was constructed, and the circles indicate sites were *COX I* data was collected. The asterisks indicate cases where *COX I* data was downloaded. **D**. PCA with Pool-seq data from all populations. The colors represent populations. Pacific Canada (WCAN; pink), Maine (ME; blue), Rhode Island (RI; yellow), Iceland (ICE; dark green), Norway (NOR; purple), United Kingdom (UK; light green). **D-inset**. Distribution of mitochondrial haplotypes across all populations. The names *a, b* (including *b*_RI_ and *b*_SC_), and *c* represent common mtDNA haplotypes observed in populations. **E**. Nucleotide diversity (log_10_ *π*) for all nuclear genes across all populations. **F**. Tajima’s *D* for all nuclear genes across all populations. The dashed vertical line marks 0, the expected value under a neutral model. The y-axis in E and F show the density of observations. **G**. Demographic reconstruction for North Atlantic individuals showing demographic changes from 2 mya to 200 kya. **I**. demographic changes in British and Norwegian individuals. **H**. North Pacific individual showing demographic changes from 2 mya to 200 kya. **J**. Plot of recent (today – 250 kya) demographic changes in the North American and Icelandic individuals.

## Results

### Standing variation across oceans

Our pool-seq panels discovered ∼3M high quality single nucleotide polymorphisms (**SNPs**) across populations at common allele frequencies (>5%). When linkage is removed at 500 bp, the SNP panel thins to ∼690,000. Principal component analysis (PCA), on the LD-thinned SNPs, shows that variation is strongly subdivided by ocean basins (**Fig. 1D**). PC 1 captures 74% of the variation, and partitions populations across basins. PC 2 (8.5% var.) partitions Atlantic populations into 2 discrete east-west clusters. The western cluster contains ME, RI, ICE, and the eastern cluster contains UK and NOR. These clusters are supported by the abundance of mtDNA haplotypes within and between ocean basins (**Fig. 1D inset; Table S1**)(Wares and Cunningham 2001; Flight, et al. 2012; Nunez, et al. 2018). The large divergence between oceans is also captured in levels of nucleotide diversity (***π***; a metric of standing genetic variation). Surprisingly, North Atlantic populations harbor more genetic variation (*π* = 1.05%) than their Pacific, ancestral, conspecifics (*π* = 0.55%; **Fig. 1E; Fig. S1**). We also estimated the Tajimas’ *D* statistic (***D***), a measure of the excess (*D*<0), or deficit (*D*>0), of rare alleles in populations. These data indicate that all North Atlantic populations, especially NOR, have negatively skewed genome-wide values of *D* (**Figs. 1E, S2**).

### Historical phylogeography and structure

We reconstructed changes of historical effective population sizes (***N*_e_**) with the multiple sequentially Markovian coalescent model (MSMC) using individual whole genomes (Schiffels and Durbin 2014). Our results provide evidence for different phylogeographic trajectories in response to the events of the glaciations (**Figs. 1G, 1H**). For instance, the Eastern Cluster and the Western Cluster populations shared a common demography throughout the Pleistocene (**Fig. 1G**) but diverged in recent geological time. Namely, Eastern populations (especially NOR) experienced striking increases in *N*_*e*_ in the recent past (**Fig. 1I**), likely following the asynchronous deglaciation of the Fennoscandian ice sheet (Ruddiman and Mcintyre 1981; Patton, et al. 2017). Western populations, on the other hand, experienced a demographic contraction which started during the last glacial period and ended during the last glacial maxima (∼20 kya; **Fig. 1J**)(Brochmann, et al. 2003; Maggs, et al. 2008; Flight, et al. 2012).

We estimated gene flow by computing *f*_3_ statistics (Reich, et al. 2009) for all possible combinations of target, source 1, and source 2 populations, using individual whole genomes (**Fig. S3; Table S2**). Our analysis finds no evidence of recent gene flow across oceans. This result is supported by two additional lines of evidence. First, a mtDNA molecular clock analysis (Drummond, et al. 2002), which suggests that Pacific and Atlantic populations have not exchanged migrants in nearly 2 million years (**SI Appendix 3**). And second, estimates of genetic differentiation (i.e., ***F*_ST_**) which reveal large amounts of genome-wide divergence (**Fig. S4**), and foreshadows the onset of allopatric speciation across oceans. Within the North Atlantic, *F*_ST_ is low (likely due to shared demography until the glacial maximum) and the *f*_3_ analysis suggest that admixture is pervasive (**Fig. S3, Table S2**). These findings are supported by additional ABBA-BABA tests for gene tree heterogeneity (Green, et al. 2010) (see **SI Appendix 4**). Overall, these findings present three important points. First, they exemplify the complex demography that underlie standing variation in natural populations. Second, they confirm that barnacles harbor high levels of genetic variation genome-wide. And third, they reveal the pervasiveness of gene flow and shared variation within ocean basins, where environmental heterogeneity is extensive across “micro” (1-3 meter) and “meso” (1-10 kilometer) scales. These conditions provide the environmental context for ecological load at the genomic scale.

### Balancing selection in barnacles

Balancing selection is expected to produce molecular and phylogenetic footprints not consistent with neutrality (Fijarczyk and Babik 2015). Molecular footprints include: enrichment of old alleles (e.g., trans-species polymorphisms; **TSPs**), elevated genetic variation (high *π*), deficit of rare alleles (*D* > 0), excess SNPs at medium allele frequencies, reduced divergence around the balanced locus (low *F*_ST_), as well as the accumulation of non-synonymous variation in the vicinity of balanced polymorphisms, a phenomenon known as sheltered load (Uyenoyama 2005). Likewise, balancing selection will produce a phylogenetic signal composed of diverged clades, corresponding to the balanced haplotypes. Deeply diverged clades will occur when balancing selection has maintained variation over long evolutionary times (i.e., ancestral balancing selection(Fijarczyk and Babik 2015)). A joint analysis of our Pacific, Atlantic, and outgroup (*S. cariosus*) datasets reveal 11,917 cosmopolitan SNPs (i.e., SNPs that segregate in both oceans) which are also TSPs (**Dataset S1**). TSPs, genome-wide, occur in 0.14% coding regions, 0.21% in introns, 0.02 % in promoters, 0.01% in 5’UTRs, and < 0.01% in 3’UTRs. The remainder TSPs occur in 0.09% of intergenic regions. An enrichment analysis which compares the abundance of TSPs, of each genomic class, relative to all discovered SNPs, reveals that TSPs are significantly over-enriched in coding loci (**Fig. 2A**), and 4,415 segregate at high frequencies in all populations (TSPs with heterozygosity [***H*_E_**] > 0.30; **Fig. S5**). These patterns of variation could be the result of neutral processes such as recurrent mutation (homoplasy) across all populations of either species. However, the enrichment of cosmopolitan, nonsynonymous, TSPs at common frequencies is not consistent with neutrality. Under a model of strict neutrality, segregating mutations are eventually lost in populations after speciation (Clark 1997). Moreover, coding regions are subjected to purifying selection which removes deleterious and mildly deleterious nonsynonymous variants (Hartl and Clark 1997).

**Figure 2.**
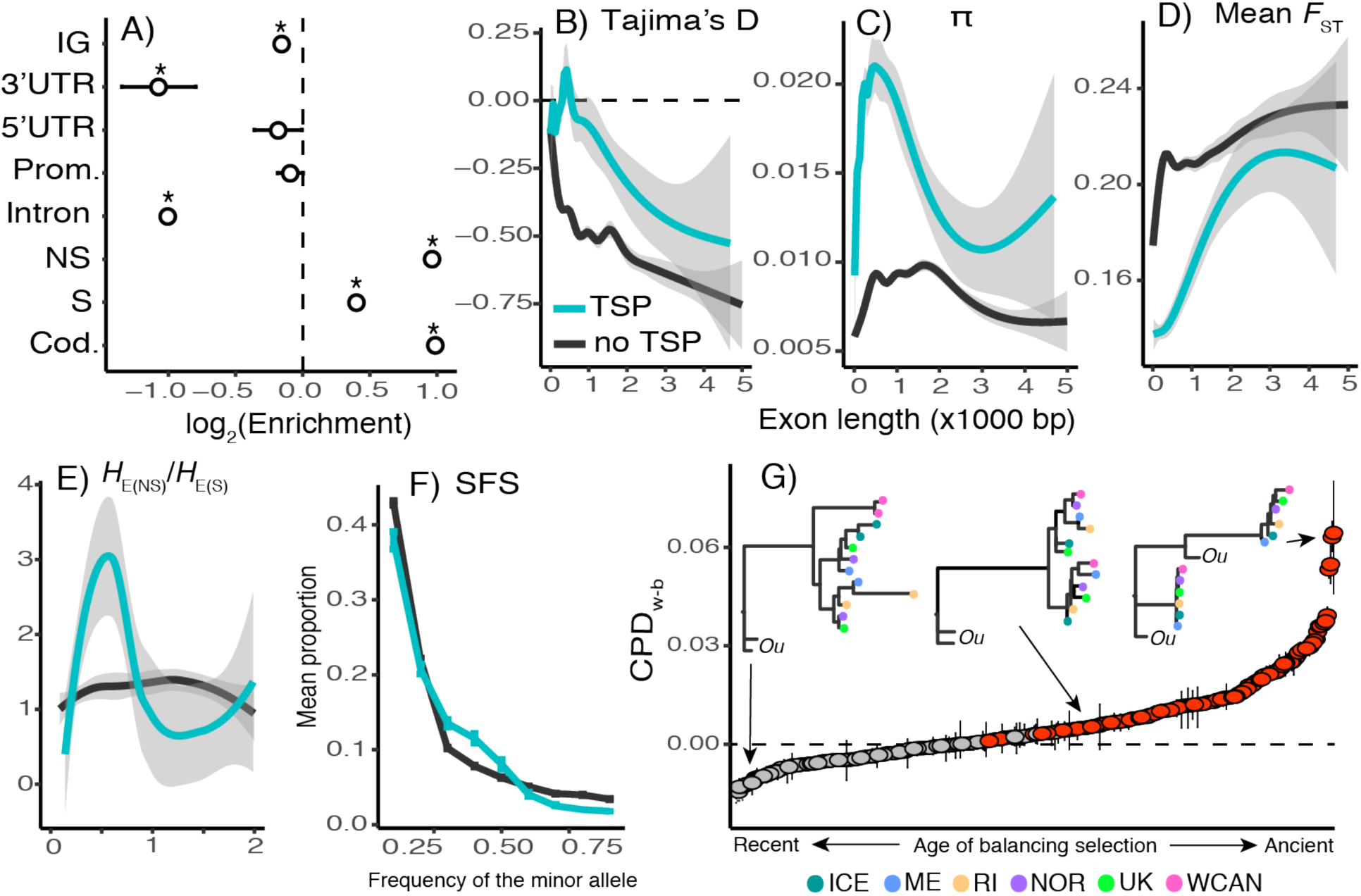
Evidence for balancing selection across the genome. **A**. Enrichment analysis of TSPs across the genome of *S. balanoides* based on all populations studied. The asterisks symbols represent statistical significance. Abbreviations: promoters (Prom.), nonsynonymous loci (NS), synonymous loci (S), coding loci (Cod.). **B**. Plot of Tajima’s *D* (as a function of length) of exons bearing TSPs versus all other exons not bearing TSPs. **C**. Same as B but for nucleotide diversity (*π*). **D**. Same as B but for mean *F*_ST_. **E**. Same as B but for the ratio of nonsynonymous heterozygosity to synonymous heterozygosity. **F**. Site frequency spectrum for whole genes with TSPs vs other genes. Vertical bars are 95% confidence intervals. **G**. Candidate genes under balancing selection ranked according to their CPD_w-b_ values (interquartile ranges shown as error bars). Red values indicate statistical significance. Horizontal dashed line indicates CPD_w-b_ = 0. Three example allele tree topologies are shown. The sister taxon, *S. cariosus*, is shown as “*Ou*” (for outgroup). The x-axis for B, C, D, and E is exon length (x 1000 bp).

We compared patterns of genetic variation in exons bearing TSPs and other exons. When accounting for exon length, we observe consistently elevated values of *D* and *π* for TSP-bearing exons relative to other exons (**Figs. 2B and 2C**; **S6**). Except for the ME vs. RI comparison (**Fig. S7**), TSP-bearing exons have consistently low *F*_ST_ values (**Fig. 2D**). To quantify sheltered load, we compared the ratio of *H*_E_ values at nonsynonymous (**NS**) and synonymous (**S**) mutations in TSP-bearing and other exons. Our results show that medium sized TSP-bearing exons (∼500 bp) harbor an excess of non-synonymous NS *H*_E_ (**Fig. 2E**). Notably, we observed that differences between TSP-bearing and other exons become less apparent as exons get longer. This regionalization of the signal occurs due to the small linkage blocks in the species (Nunez, et al. 2020). We observe 1,107 TSPs that cause nonsynonymous changes and occur in 312 genes with high confidence annotations (4%; **Dataset S2**). Consistent with our expectation of balancing selection, site frequency spectrum (**SFS**) analyses show that these 312 genes harbor an excess of SNPs at medium allele frequencies relative to other annotated genes (**Fig. 2F**).

### Age of balanced polymorphisms

To determine the age of balanced polymorphisms, we ran topological tests on the allele trees for each TSP region across the 312 candidate genes. We built trees using phased haplotypes for each TSP-bearing region for all single individual genomes. We used these allele trees to compute the cophenetic distance (**CPD**) between tips. We classified allele trees as having or lacking highly diverged alleles based on the relative mean CPD between haplotypes from the same population vs. from different populations (CPD_w-b_; **see supplementary methods**). The analysis reveals that of the 312 allele trees, 150 carry a significant signature of ancestral balancing selection (CDP_w-b_ > 0, Bonferroni *P* < 1×10^−9^; **Fig. 2G; Dataset S2**). This suggests maintenance of diverged haplotypes for more than 2 million years, with extreme cases in which haplotypes are shared across species (8-1o million years)(Perez-Losada, et al. 2008; Herrera, et al. 2015). The remaining genes with CDP_w-b_ < 0 may either represent cases where the balanced alleles are younger, or oversampling of homozygous individuals for any given marker.

### Targets of selection

We partitioned our dataset among genes with positive and negative CPD_w-d_ allele trees and conducted gene ontology (GO) enrichment analyses. The 150 genes with positive CPD_w-d_ trees show enrichment for terms related to “ion channel regulation”, including genes involved in environmental sensing, and circadian rhythm regulation (**Table S3**). We show examples for 3 candidate genes under ancestral balancing selection involved in environmental sensing: 1) the painless gene (*Pain*; g1606; **Fig. 3A**), which is involved in nociception (i.e., pain reception), as well as detection of heat and mechanical stimuli (Tracey, et al. 2003; Xu, et al. 2006); 2) the Pyrexia gene (*Pyx*; g3472; **Fig. 3B**), which is involved in negative geotaxis, and responses to heat (Lee, et al. 2005); and 3) the shaker cognate w gene (*Shaw*; g3310; **Fig. 3C**), which is involved in regulation of circadian rhythm (Hodge and Stanewsky 2008; Buhl, et al. 2016). These three examples showcase canonical footprints of balancing selection around the TSP, concomitant with a bimodal allele tree. Among genes with negative CPD_w-d_ we observe enriched functions for “anatomical structure formation” including genes coding for motor proteins and muscle genes (**Table S4**). In all cases, we used RNA-seq data from ME individuals to confirm that these loci are expressed in adult barnacles.

**Figure 3.**
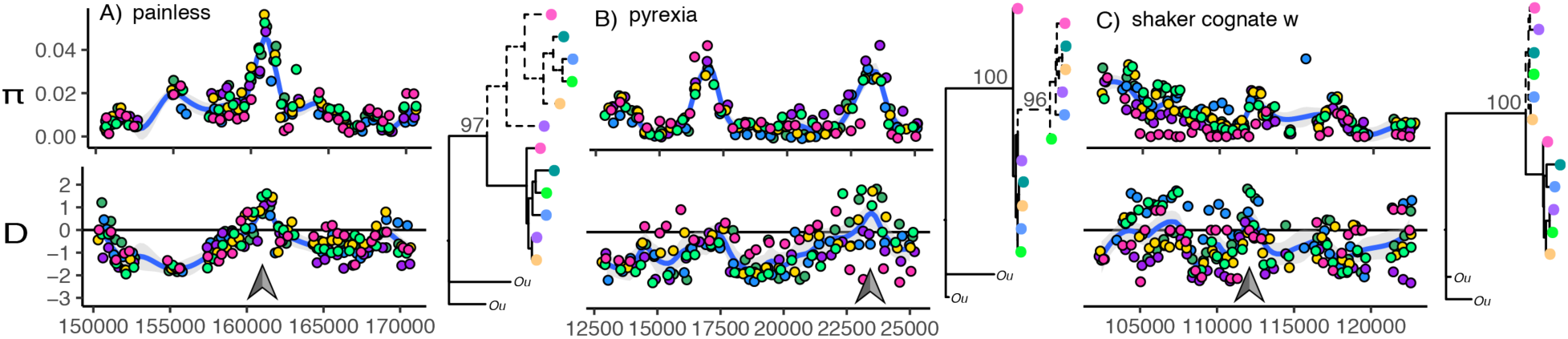
Balancing selection on ecologically important genes. We present patterns of genetic variation (*π* and *D* estimated from pool-seq data, and allele tree topologies estimated from single individuals) for 3 example genes: A) painless (*Pain*), B) pyrexia (*Pyx*), C) shaker cognate w (*Shaw*). Grey arrows show regions that contain TSPs. In Tajima’s *D* panels, the horizontal line marks the *D* = 0 point. For all trees, the sister taxon, *S. cariosus*, is shown as “*Ou*”. The colors represent populations. WCAN (pink), ME (blue), RI (yellow), ICE (dark green), NOR (purple), UK (light green). The x-axis shows base pair position within scaffolds.

## Discussion

In intertidal barnacles, the dichotomy of strong adult selection and high offspring dispersal means that any allele that is beneficial to parental fitness in one generation may be neutral or deleterious in the next (Gillespie 1973). This leads to a fundamental question in evolutionary biology: how are adaptations maintained in the face of extreme ecological variability? In this paper, we provide evidence that balancing selection is widespread across the barnacle genome, with 4% of annotated genes harboring functional balanced polymorphisms. Notably, these polymorphisms occur in genes with important functions for life in variable environments, and many have been maintained for at least 2 million years despite a complex phylogeographic history (Wares and Cunningham 2001; Flight and Rand 2012). Naturally, the heterogeneous nature of the rocky intertidal imposes a segregation ‘cost’ for these balanced polymorphisms, as they occur in individuals that, due to high dispersal, recruit in sub-optimal habitats for any given genetic makeup. This ecological load, defined as 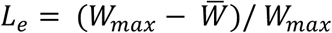 (where 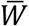 is mean fitness, and *W*_*max*_ is optimal fitness, across all habitats), will be substantial, as demonstrated by comprehensive recruitment studies in natural habitats (Bertness 1989; Bertness, et al. 1992; Pineda, et al. 2006). For example, at initial settlement, barnacle density can be as high as 76 individuals per cm^2^, but at maturity, it can be as low as 0.15 individuals per cm^2^ (0.2% survival)(Pineda, et al. 2006). This mass mortality is habitat- and genotype-dependent (Schmidt and Rand 2001). This is the type of ‘fitness cost’ envisioned in the Levene model of balancing selection (Levene 1953). As such, our data suggests that the problem of ecological load is a defining condition of the barnacle life cycle. And, more generally, it argues in favor of balancing selection, via marginal overdominance, as the fundamental process underlying maintenance of adaptation in variable environments.

### Is pervasive balancing selection plausible in nature?

Under classical models of population genetics, when loci are considered to be independent of each other, the additive effects of widespread balanced polymorphism results in unbearable amounts of fitness variance and genetic death (Kimura and Crow 1964; Lewontin and Hubby 1966). However, if balanced loci have interactive effects (e.g., epistasis), multiple polymorphisms could be maintained with minimum effects on the distribution of fitness variance (King 1967; Milkman 1967; Sved, et al. 1967; Wittmann, et al. 2017). Based on this theoretical framework, multiple models have been developed to describe the conditions that favor the long-term maintenance of functional variation in spatially varying environments (Gillespie 1973; Hedrick, et al. 1976). Moreover, polymorphisms will be less likely to be lost if there is a large number of ecological niches available, if there is migration among niches, and if individuals are proactive in choosing niches where their fitness is maximized (Hedrick, et al. 1976). We argue that barnacles satisfy these conditions to some degree.

First, while it is useful to summarize intertidal heterogeneity in the form of discrete microhabitats (Schmidt, et al. 2000), individual barnacles experience the rocky shore as a complex tapestry of interactive stressors at three spatial levels. At microhabitats scales, the upper and lower tidal zones pose diametrically different ecological challenges in terms of food availability, competition, predation, and risk of desiccation (Bertness, et al. 1991; Schmidt and Rand 1999, 2001). At mesohabitat scales, open coasts vs. sheltered estuaries vary in their exposure to wave action, upwelling dynamics, and biotic interactions (Sanford and Menge 2001; Dufresne, et al. 2002; Veliz, et al. 2004). These, in turn, modify micro-level stressors. Lastly, at macrohabitat scales, topological differences across shores and latitudinal variations in tidal range produce a mosaic of thermal stress along continents (Helmuth, et al. 2002). Consequentially, what selection pressures are more important for any given barnacle will emerge from the interactions among these stress gradients. This complex landscape of selection has been captured in studies of the barnacle *Mpi* gene. Accordingly, the locus is under selection at micro-levels in the Gulf of Maine (Schmidt and Rand 1999; Schmidt, et al. 2000), at meso-levels in the gulf of St. Lawrence (Canada)(Dufresne, et al. 2002; Veliz, et al. 2004), yet it shows tepid signs of selection in the Narragansett Bay (Rhode Island)(Rand, et al. 2002; Nunez, et al. 2020). Similar complexity has also been captured in temperate populations of *Drosophila*. In these, idiosyncratic weather effects can alter the dynamics of seasonal adaptation (Bergland, et al. 2014; Machado, et al. 2019). Second, the high dispersal capacity of the larval stage ensures constant migration between these niches across generations. Finally, barnacles also have the ability to choose preferred substrates during settlement. This occurs during the spring when barnacle larvae extensively survey microhabitats for biological, chemical and physical cues produced by previous settlers before making final commitments of where to settle (Bertness, et al. 1992). Unfortunately for the barnacle, this capacity for substrate choice does not mitigate mass mortality during late summer, which leads to strong selection for particular genotypes (Schmidt and Rand 2001). Nevertheless, these behaviors may constitute a form of adaptive plasticity, helping barnacles choose habitats where their fitness may be marginally improved. Overall, this suggests that the barnacle’s life history is conducive to the maintenance of balanced polymorphisms.

### What variation is under selection?

Our analyses indicate that 4% (312) of all annotated genes are experiencing some form of balancing selection across the entire range of the species. This number of genes harboring ancestral polymorphisms is similar to that observed in *Arabidopsis thaliana* and its close relative *Capsella rubella* (433 genes)(Wu, et al. 2017). Similar to *Semibalanus*, these plants diverged ∼8 mya, and their natural populations experience high levels of ecological heterogeneity (Bakker, et al. 2006). We must acknowledge that our number may be an underestimation driven by the nascent state of the genomic tools in *Semibalanus*. Future genome assemblies, combined with improved annotations, will undoubtedly yield a more complete picture of functional variation in the species. In addition, it will allow for a more comprehensive characterization of selection in structural variants and regulatory loci, which have been shown to be fundamental in the evolution of complex phenotypes (Wray 2007; Faria, et al. 2019). Despite these limitations, our analysis recovered a large number of genes involved in key functions for life in variable environments. These will be subjects of future validation studies. For instance, the general enrichment for ion channel genes suggests selection related to osmotic regulation (Sundell, et al. 2019). This hypothesis is highly plausible given that intertidal ecosystems experience strong salinity fluctuations, repeatedly exposing barnacles to osmotic challenges at all spatial scales. In addition, we observe targets of selection involved in environmental sensing loci (e.g., *pain, pyx*, and *shaw;* **Fig. 3**). Similar to osmotic regulation, selection on these genes is entirely plausible given the inherent variability of intertidal habitats. An important hypothesis from the allozyme era is the idea that balancing selection would target genes at the node of metabolic fluxes (Eanes 1999; Watt and Dean 2000). In such cases, balanced variation would provide biochemical flexibility to cope with environmental heterogeneity. In the same vein, we hypothesize that balancing selection may act more often on “sensor genes” which control plastic responses to ecological variation. Testing this hypothesis is beyond the scope of this paper and would require the use of allele-specific differential expression experiments in barnacles.

### Complex demography and speciation

Our demographic analyses provide clues about how historical events affected genetic variation in barnacle populations. In the Atlantic, our evidence suggests a shared demography throughout the Pleistocene, and that the modern Eastern and Western clusters formed in response to recent events of last glacial cycle. These findings highlight that the low *F*_ST_ values observed within the basins arise due to shared ancestry. Moreover, they also suggest that population structure persists in the presence of gene flow. As such, while larvae have the capacity to disperse for hundreds of kilometers, ocean currents (Nunez, et al. 2018) and different estuarine flushing times (Brown, et al. 2001) allow regions to retain some level of geographical structuring (Johannesson, et al. 2018; Nunez, et al. 2018). Comparisons between oceans reveal a stark pattern of genome wide divergence. This pattern is driven by the separation of Pacific and Atlantic populations following the events of the trans-Arctic interchange (Vermeij 1991). Accordingly, the negative levels of *D* in the north Atlantic may reflect the effect of bottlenecks during the trans-Arctic interchange. Notably, the high levels of *π* in the Atlantic is not concordant with predictions of common colonization models in which variation of the younger population is a subset of the ancestral population (Maggs, et al. 2008). We hypothesize this could be the result of ancient admixture due to repeated trans-Arctic invasions from the Pacific (Väinölä 2003). We recognize that ancestral admixture could generate artificial signatures of balancing selection via the mixing of highly differentiated haplotypes. However, such an occurrence would affect most genes in the genome. Our evidence shows that the signatures of balancing selection are highly localized in TSP-regions. For example, while *D* is elevated in TSP-regions, it is negatively skewed genome-wide. Our data does not support recent gene flow between ocean basins. As such, after 2 million years of separation, neutral divergence appears to be driving Atlantic and Pacific populations to speciate in allopatry. A closer look to this hypothesis will require crossing individuals from both basins, and surveying offspring fitness and viability. More salient, however, is the observation of shared haplotypes between oceans in our candidate genes for balancing selection. In light of such strong background divergence, this provides evidence that balancing selection on most of these genes is strong, and that polymorphisms have been maintained for long periods of time.

## Materials & Methods

### Barnacle Collections

Barnacle samples were collected from Damariscotta (Maine, United States; ME), Jamestown (Rhode Island, United States, RI), Calvert Island (British Columbia, Canada; WCAN), Reykjavik (Iceland; ICE), Porthcawl (Wales, United Kingdom; UK), and Norddal (Norway; NOR). Additional samples were collected in Bergen (Norway), Tórshavn (Faroe Island), and Tjärnö (Sweden). For all samples, species identities were confirmed using Sanger sequencing of the mtDNA COX I region(Bucklin, et al. 2011). For the WCAN, RI, ME, ICE, UK, and NOR population we collected a single individual for DNA-seq, and a group of 20-40 individuals for pool-seq (**SI Appendix 2**). RNA-seq was done on four individuals from Maine. DNA-seq was done on a single individual from the sister taxa *S. cariosus*. DNA/RNA was extracted using Qiagen DNeasy/RNeasy kits. All pools and single individuals were sequenced in their own lanes of an Illumina machine by GENEWIZ LLC using 2×150 paired-end configuration.

### Mapping datasets to the genome

Samples were mapped to a genome assembled *de novo* for the species (Sbal3.1; NCBI GenBank: VOPJ00000000; **SI Appendix 1**). The genome was assembled using a hybrid approach which combines PacBio reads and Illumina reads using DBG2OLC(Ye, et al. 2016) and Redundans(Pryszcz and Gabaldon 2016). Gene models were constructed using an *ab initio* method, AUGUSTUS(Stanke and Waack 2003), informed by evidence from the RNA-seq. A gene feature file (GFF) is available as **Dataset S4**. The model used for gene prediction was trained in *Drosophila melanogaster*. Genes were annotated by pairwise blast against the *Drosophila melanogaster* genome (Dmel6; NCBI GenBank: GCA_000001215.4). All annotations are available as **Dataset S5**. DNA reads from all populations were mapped to Sbal3.1 using bwa mem(Li 2013). RNA reads were mapped using HiSat2(Kim, et al. 2015). SNPs were called using the samtools pipeline(Li, et al. 2009).

### Genome analyses

Estimates of *π* and *D* were done using the popoolation-1 suite(Kofler, Orozco-terWengel, et al. 2011). Estimations of allele frequencies and *F*_ST_ were done using the popoolation-2 suite(Kofler, Pandey, et al. 2011). Demographic reconstructions were done using MSMC(Schiffels and Durbin 2014). The *f*_3_ statistics were estimated using treemix(Pickrell and Pritchard 2012). Bayesian molecular clock analyses were done in BEAST2(Bouckaert, et al. 2014). ABBA/BABA statistics were calculated in *Dsuite*(Malinsky, et al. 2020). Phylogenetic inferences were done in iQtree(Chernomor, et al. 2016). GO enrichment analysis was done using GOrilla(Eden, et al. 2009) and GO terms inferred from our *Drosophila* annotation. The enrichment was assessed by comparing 2 genes list. The first composed of the genes of interest (i.e., the gene targets), the second one by all the genes annotated in Sbal3.1 (i.e., the gene universe). A detailed description of our analyses can be found in the supplementary methods section, as well as in GitHub: **https://github.com/Jcbnunez/BarnacleEcoGenomics**.

## Acknowledgements

All the authors would like to acknowledge the Centre for Marine Evolutionary Biology (CeMEB) at the University of Gothenburg which organized the Marine Evolution 2018 meeting in which most of the authors met and started this collaborative work. In addition, we acknowledge S. Ramachandran, E. Huerta-Sanchez, D. Sax, D. R. Gaddes, and R. E. F. Gordon, for their support and helpful insights. To E. Sanford for providing the sample of *S. cariosus*. To M. D. Rand and family for collecting barnacles in Nordall, Norway. To C. Harley for providing the samples from British Columbia. We thank the Natural Environment Research Council, the European Research Council, the Swedish Research Councils VR and Formas (Linnaeus grant to CeMEB), and *SciLife* Laboratory. This research was conducted using computational resources and services at the Center for Computation and Visualization, Brown University.

## Data deposition

Data used in this paper are available in the National Center for Biotechnology Information (NCBI), https://www.ncbi.nlm.nih.gov. Raw reads were deposited under submission id: SUB6188969. SRAs are as follows: DNAseq datasets: SRR10011798, SRR10011802, SRR10011804, SRR10011805, SRR10011807–SRR10011810, SRR10011812–SRR10011814, SRR10011819, SRR10011825; PacBio dataset: SRR10011818; RNAseq datasets: SRR10011820-SRR10011823. MtDNA sequences for the COX I genes can be acceded form the following GeneBank accessions MG925538– MG925662, MG928281–MG928323, and MT329074–MT329592. Whole mtDNAs were deposited under accessions MG010647, MG010648, MG010649, MT528636, MT528637. The barnacle genome (Sbal3.1) is available at NCBI (accession no. VOPJ00000000). A GitHub repository with code as well as with the supplementary datasets S1, S2, S3, S4, and S5, can be found at **https://github.com/Jcbnunez/BarnacleEcoGenomics**.

## References

Bakker EG, Stahl EA, Toomajian C, Nordborg M, Kreitman M, Bergelson J. 2006. Distribution of genetic variation within and among local populations of Arabidopsis thaliana over its species range. Mol Ecol 15:1405–1418.

Bergland AO, Behrman EL, O’Brien KR, Schmidt PS, Petrov DA. 2014. Genomic Evidence of Rapid and Stable Adaptive Oscillations over Seasonal Time Scales in Drosophila. PLoS Genet 10:e1004775.

Bertness MD. 1989. Intraspecific Competition and Facilitation in a Northern Acorn Barnacle Population. Ecology 70:257–268.

Bertness MD, Gaines SD, Bermudez D, Sanford E. 1991. Extreme spatial variation in the growth and reproductive output of the acorn barnacle Semibalanus balanoides. Marine Ecology Progress Series 75:91–100.

Bertness MD, Gaines SD, Stephens EG, Yund PO. 1992. Components of recruitment in populations of the acorn barnacle Semibalanus balanoides (Linnaeus). Journal of Experimental Marine Biology and Ecology 156:199–215.

Botero CA, Weissing FJ, Wright J, Rubenstein DR. 2015. Evolutionary tipping points in the capacity to adapt to environmental change. Proc Natl Acad Sci U S A 112:184–189.

Bouckaert R, Heled J, Kuhnert D, Vaughan T, Wu CH, Xie D, Suchard MA, Rambaut A, Drummond AJ. 2014. BEAST 2: a software platform for Bayesian evolutionary analysis. PLoS Comput Biol 10:e1003537.

Brochmann C, Gabrielsen TM, Nordal I, Landvik JY, Elven R. 2003. Glacial survival or tabula rasa? The history of North Atlantic biota revisited. Taxon 52.

Brown AF, Kann LM, Rand DM. 2001. Gene flow versus local adaptation in the northern acorn barnacle, Semibalanus balanoides: insights from mitochondrial DNA variation. Evolution 55:1972–1979.

Bucklin A, Steinke D, Blanco-Bercial L. 2011. DNA Barcoding of Marine Metazoa. Annu. Rev. Mar. Sci. 3:471–508.

Buhl E, Bradlaugh A, Ogueta M, Chen KF, Stanewsky R, Hodge JJ. 2016. Quasimodo mediates daily and acute light effects on Drosophila clock neuron excitability. Proc Natl Acad Sci U S A 113:13486–13491.

Chernomor O, von Haeseler A, Minh BQ. 2016. Terrace Aware Data Structure for Phylogenomic Inference from Supermatrices. Syst Biol 65:997–1008.

Clark AG. 1997. Neutral behavior of shared polymorphism. Proc Natl Acad Sci U S A 94:7730–7734.

Drummond AJ, Nicholls GK, Rodrigo AG, Solomon W. 2002. Estimating mutation parameters, population history and genealogy simultaneously from temporally spaced sequence data. Genetics 161:1307–1320.

Dufresne F, Bourget E, Bernatchez L. 2002. Differential patterns of spatial divergence in microsatellite and allozyme alleles: further evidence for locus-specific selection in the acorn barnacle, Semibalanus balanoides? Molecular Ecology 11:113–123.

Eanes WF. 1999. Analysis of Selection on Enzyme Polymorphisms. Annual Review of Ecology and Systematics 30:301–326.

Eden E, Navon R, Steinfeld I, Lipson D, Yakhini Z. 2009. GOrilla: a tool for discovery and visualization of enriched GO terms in ranked gene lists. Bmc Bioinformatics 10:48.

Faria R, Johannesson K, Butlin RK, Westram AM. 2019. Evolving Inversions. Trends Ecol Evol 34:239–248.

Fijarczyk A, Babik W. 2015. Detecting balancing selection in genomes: limits and prospects. Mol Ecol 24:3529–3545.

Flight PA, O’Brien MA, Schmidt PS, Rand DM. 2012. Genetic Structure and the North American Postglacial Expansion of the Barnacle, Semibalanus balanoides. Journal of Heredity 103:153–165.

Flight PA, Rand DM. 2012. Genetic variation in the acorn barnacle from allozymes to population genomics. Integrative and Comparative Biology 52:418–429.

Flowerdew MW. 1983. Electrophoretic Investigation of Populations of the Cirripede Balanus-Balanoides (L) around the North-Atlantic Seaboard. Crustaceana 45:260–278.

Gillespie J. 1973. Polymorphism in random environments. Theoretical population biology 4:193–195.

Green RE, Krause J, Briggs AW, Maricic T, Stenzel U, Kircher M, Patterson N, Li H, Zhai W, Fritz MH, et al. 2010. A draft sequence of the Neandertal genome. Science 328:710–722.

Hartl DL, Clark AG. 1997. Principles of population genetics: Sunderland: Sinauer associates.

Hedrick PW. 2006. Genetic polymorphism in heterogeneous environments: The age of genomics. Annual Review of Ecology Evolution and Systematics 37:67–93.

Hedrick PW, Ginevan ME, Ewing. EP. 1976. Genetic polymorphism in heterogeneous environments. Annual Review of Ecology and Systematics 7.

Helmuth B, Harley CD, Halpin PM, O’Donnell M, Hofmann GE, Blanchette CA. 2002. Climate change and latitudinal patterns of intertidal thermal stress. Science 298:1015–1017.

Herrera S, Watanabe H, Shank TM. 2015. Evolutionary and biogeographical patterns of barnacles from deep-sea hydrothermal vents. Mol Ecol 24:673–689.

Hodge JJ, Stanewsky R. 2008. Function of the Shaw potassium channel within the Drosophila circadian clock. PLoS ONE 3:e2274.

Johannesson K, Ring AK, Johannesson KB, Renborg E, Jonsson PR, Havenhand JN. 2018. Oceanographic barriers to gene flow promote genetic subdivision of the tunicate Ciona intestinalis in a North Sea archipelago. Mar Biol 165:126.

Jones SJ, Southward AJ, Wethey DS. 2012. Climate change and historical biogeography of the barnacle Semibalanus balanoides. Global Ecology and Biogeography 21:716–724.

Kim D, Langmead B, Salzberg SL. 2015. HISAT: a fast spliced aligner with low memory requirements. Nat Methods 12:357–360.

Kimura M, Crow J. 1964. The number of alleles that can be maintained in a finite population. Genetics 49:725–738.

King JL. 1967. Continuously distributed factors affecting fitness. Genetics 55:483–492.

Kofler R, Orozco-terWengel P, De Maio N, Pandey RV, Nolte V, Futschik A, Kosiol C, Schlotterer C. 2011. PoPoolation: a toolbox for population genetic analysis of next generation sequencing data from pooled individuals. PLoS ONE 6:e15925.

Kofler R, Pandey RV, Schlotterer C. 2011. PoPoolation2: identifying differentiation between populations using sequencing of pooled DNA samples (Pool-Seq). Bioinformatics 27:3435–3436.

Lee Y, Lee Y, Lee J, Bang S, Hyun S, Kang J, Hong ST, Bae E, Kaang BK, Kim J. 2005. Pyrexia is a new thermal transient receptor potential channel endowing tolerance to high temperatures in Drosophila melanogaster. Nat Genet 37:305–310.

Levene H. 1953. Genetic Equilibrium When More Than One Ecological Niche Is Available. American Naturalist 87:331–333.

Lewontin RC, Hubby JL. 1966. A molecular approach to the study of genic heterozygosity in natural populations. II. Amount of variation and degree of heterozygosity in natural populations of Drosophila pseudoobscura. Genetics 54:595–609.

Li H. 2013. Aligning sequence reads, clone sequences and assembly contigs with BWA-MEM. 1303.3997.

Li H, Handsaker B, Wysoker A, Fennell T, Ruan J, Homer N, Marth G, Abecasis G, Durbin R, Genome Project Data Processing S. 2009. The Sequence Alignment/Map format and SAMtools. Bioinformatics 25:2078–2079.

Machado HE, Bergland AO, Taylor R, Tilk S, Behrman E, Dyer K, Fabian DK, Flatt T, González J, Karasov TL, et al. 2019. Broad geographic sampling reveals predictable and pervasive seasonal adaptation in Drosophila. bioRxiv (unpublished data) https://www.biorxiv.org/content/biorxiv/early/2019/10/11/337543.full.pdf - last accessed December 12, 2019.

Mackay TF, Richards S, Stone EA, Barbadilla A, Ayroles JF, Zhu D, Casillas S, Han Y, Magwire MM, Cridland JM, et al. 2012. The Drosophila melanogaster Genetic Reference Panel. Nature 482:173–178.

Maggs CA, Castilho R, Foltz D, Henzler C, Jolly MT, Kelly J, Olsen J, Perez KE, Stam W, Väinölä R, et al. 2008. Evaluating Signatures of Glacial Refugia for North Atlantic Benthic Marine Taxa. Ecology 89:S108–S122.

Malinsky M, Matschiner M, Svardal H. 2020. Dsuite - fast D-statistics and related admixture evidence from VCF files. bioRxiv (unpublished data) https://www.biorxiv.org/content/10.1101/634477v2 - last accessed May 1, 2020.

Messer PW, Petrov DA. 2013. Population genomics of rapid adaptation by soft selective sweeps. Trends Ecol Evol 28:659–669.

Metz EC, Palumbi SR. 1996. Positive selection and sequence rearrangements generate extensive polymorphism in the gamete recognition protein bindin. Mol Biol Evol 13:397–406.

Milkman RD. 1967. Heterosis as a Major Cause of Heterozygosity in Nature. Genetics 55:493-&.

Nunez JCB, Elyanow RG, Ferranti DA, Rand DM. 2018. Population Genomics and Biogeography of the Northern Acorn Barnacle (Semibalanus balanoides) Using Pooled Sequencing Approaches. In: Oleksiak MF, Rajora OP, editors. Population Genomics: Marine Organisms: Springer, Cham. p. 139–168.

Nunez JCB, Flight PA, Neil KB, Rong S, Eriksson LA, Ferranti DA, Rosenblad MA, Blomberg A, Rand DM. 2020. Footprints of natural selection at the mannose-6-phosphate isomerase locus in barnacles. Proc Natl Acad Sci U S A:201918232.

Patton H, Hubbard A, Andreassen K, Auriac A, Whitehouse PL, Stroeven AP, Shackleton C, Winsborrow M, Heyman J, Hall AM. 2017. Deglaciation of the Eurasian ice sheet complex. Quaternary Science Reviews 169:148–172.

Perez-Losada M, Harp M, Hoeg JT, Achituv Y, Jones D, Watanabe H, Crandall KA. 2008. The tempo and mode of barnacle evolution. Mol Phylogenet Evol 46:328–346.

Pickrell JK, Pritchard JK. 2012. Inference of population splits and mixtures from genome-wide allele frequency data. PLoS Genet 8:e1002967.

Pineda J, Starczak V, Stueckle TA. 2006. Timing of successful settlement: demonstration of a recruitment window in the barnacle Semibalanus balanoides. Marine Ecology Progress Series 320:233–237.

Pryszcz LP, Gabaldon T. 2016. Redundans: an assembly pipeline for highly heterozygous genomes. Nucleic Acids Research 44:e113.

Rand DM, Spaeth PS, Sackton TB, Schmidt PS. 2002. Ecological Genetics of Mpi and Gpi Polymorphisms in the Acorn Barnacle and the Spatial Scale of Neutral and Non-neutral Variation. Integrative and Comparative Biology 42:825–836.

Reich D, Thangaraj K, Patterson N, Price AL, Singh L. 2009. Reconstructing Indian population history. Nature 461:489–494.

Ruddiman WF, Mcintyre A. 1981. The North-Atlantic Ocean during the Last Deglaciation. Palaeogeography Palaeoclimatology Palaeoecology 35:145–214.

Sanford E, Menge BA. 2001. Spatial and temporal variation in barnacle growth in a coastal upwelling system. Marine Ecology Progress Series 209:143–157.

Schiffels S, Durbin R. 2014. Inferring human population size and separation history from multiple genome sequences. Nat Genet 46:919–925.

Schlotterer C, Tobler R, Kofler R, Nolte V. 2014. Sequencing pools of individuals-mining genome-wide polymorphism data without big funding. Nature Reviews Genetics 15:749–763.

Schmidt PS, Bertness MD, Rand DM. 2000. Environmental heterogeneity and balancing selection in the acorn barnacle Semibalanus balanoides. Proc Biol Sci 267:379–384.

Schmidt PS, Rand DM. 2001. Adaptive maintenance of genetic polymorphism in an intertidal barnacle: habitat- and life-stage-specific survivorship of Mpi genotypes. Evolution 55:1336–1344.

Schmidt PS, Rand DM. 1999. Intertidal microhabitat and selection at Mpi: Interlocus contrasts in the northern acorn barnacle, Semibalanus balanoides. Evolution 53:135–146.

Schmidt PS, Serrão EA, Pearson GA, Riginos C, Rawson PD, Hilbish TJ, Brawley SH, Trussell GC, Carrington E, Wethey DS, et al. 2008. Ecological Genetics in the North Atlantic: Environmental Gradients and Adaptation at Specific Loci. Ecology 89:S91–S107.

Stanke M, Waack S. 2003. Gene prediction with a hidden Markov model and a new intron submodel. Bioinformatics 19:ii215–ii225.

Sundell K, Wrange AL, Jonsson PR, Blomberg A. 2019. Osmoregulation in Barnacles: An Evolutionary Perspective of Potential Mechanisms and Future Research Directions. Front Physiol 10:877.

Sved JA, Reed TE, Bodmer WF. 1967. The Number of Balanced Polymorphisms That Can Be Maintained in a Natural Population. Genetics 55:469–481.

Tracey WD, Wilson RI, Laurent G, Benzer S. 2003. painless, a Drosophila Gene Essential for Nociception. Cell 113:261–273.

Uyenoyama MK. 2005. Evolution under tight linkage to mating type. New Phytol 165:63–70.

Väinölä R. 2003. Repeated trans-Arctic invasions in littoral bivalves: molecular zoogeography of the Macoma balthica complex. Marine Biology 143:935–946.

Veliz D, Bourget E, Bernatchez L. 2004. Regional variation in the spatial scale of selection at MPI* and GPI* in the acorn barnacle Semibalanus balanoides (Crustacea). J Evol Biol 17:953–966.

Vermeij GJ. 1991. Anatomy of an invasion: the trans-Arctic interchange. Paleobiology 17:281–307.

Wares JP, Cunningham CW. 2001. Phylogeography and Historical Ecology of the North Atlantic Intertidal. Evolution 55:2455–2469.

Watt WB, Dean AM. 2000. Molecular-functional studies of adaptive genetic variation in prokaryotes and eukaryotes. Annu Rev Genet 34:593–622.

Williams G. 1966. Adaptation and Natural Selection: A Critique of Some Current Evolutionary Thought. Princeton, NJ.: Princeton University Press.

Wittmann MJ, Bergland AO, Feldman MW, Schmidt PS, Petrov DA. 2017. Seasonally fluctuating selection can maintain polymorphism at many loci via segregation lift. Proc Natl Acad Sci U S A 114:E9932–E9941.

Wray GA. 2007. The evolutionary significance of cis-regulatory mutations. Nature Reviews Genetics 8:206–216.

Wu Q, Han TS, Chen X, Chen JF, Zou YP, Li ZW, Xu YC, Guo YL. 2017. Long-term balancing selection contributes to adaptation in Arabidopsis and its relatives. Genome Biol 18:217.

Xu SY, Cang CL, Liu XF, Peng YQ, Ye YZ, Zhao ZQ, Guo AK. 2006. Thermal nociception in adult Drosophila: behavioral characterization and the role of the painless gene. Genes Brain Behav 5:602–613.

Ye C, Hill CM, Wu S, Ruan J, Ma ZS. 2016. DBG2OLC: Efficient Assembly of Large Genomes Using Long Erroneous Reads of the Third Generation Sequencing Technologies. Sci Rep 6:31900.

