## Supplemental Appendix for "Ecological load and balancing selection in circumboreal barnacles"

Supplementary material associated with this paper:

##### Figures

**Fig. S1:** Genetic variation in mtDNAs
**Fig. S2:** Additional analysis of genetic variation and Tajima's D
**Fig. S3:** *f*3 statistics graph
**Fig. S4:** Fixation index across barnacle populations
**Fig. S5:** Heterozygosity in TSPs
**Fig. S6:** Additional analysis of genetic variation and Tajima's D in TSPs
**Fig. S7:** Fixation index in TSPs

##### Tables

**Table S1:** Genetic variation in mtDNAs
**Table S2:** *f*3 statistics table
**Table S3:** Gene ontology analysis for CPD > 0
**Table S4:** Gene ontology analysis for CPD < 0

##### Appendices

**SI Appendix 1:** Details on the genome assembly
**SI Appendix 2:** Details on standing genetic variation across populations
**SI Appendix 3:** Molecular clock analysis (Pacific-Atlantic divergence)
**SI Appendix 4:** ABBA/BABA analysis
**SI Methods:** Extended methods

##### Datasets

**Dataset S1:** Dataset containing TSPs in Semibalanus
**Dataset S2:** Dataset containing Genes with TSPs and their CPDs
**Dataset S3:** Dataset containing ABBA/BABA analyses
**Dataset S4:** Gene feature file (GFF) for Sbal3.1
**Dataset S5:** Gene annotations for Sbal3.1 (based on *D. melanogaster*)

### Figure S1

Relationship between nuclear $\pi$ and mtDNA $\pi$, both $\log_{10}$ transformed. The correlation between these both values is significant at $P < 0.1$.

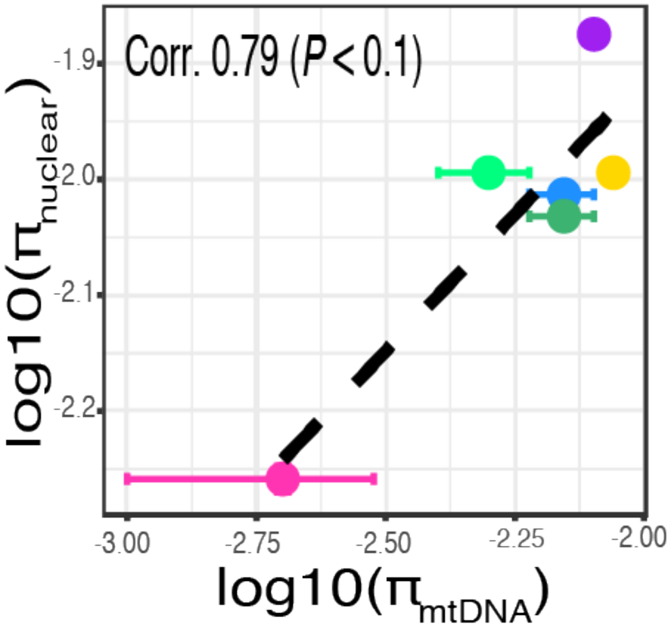

### Figure S2

Biplot of Tajima's D and $\pi$ for all genes across all populations. $\pi$ value are $\log_{10}$ transformed. The density of genes at given values is indicated by color intensity from grey to orange. The horizontal line indicates Tajima's D = 0.

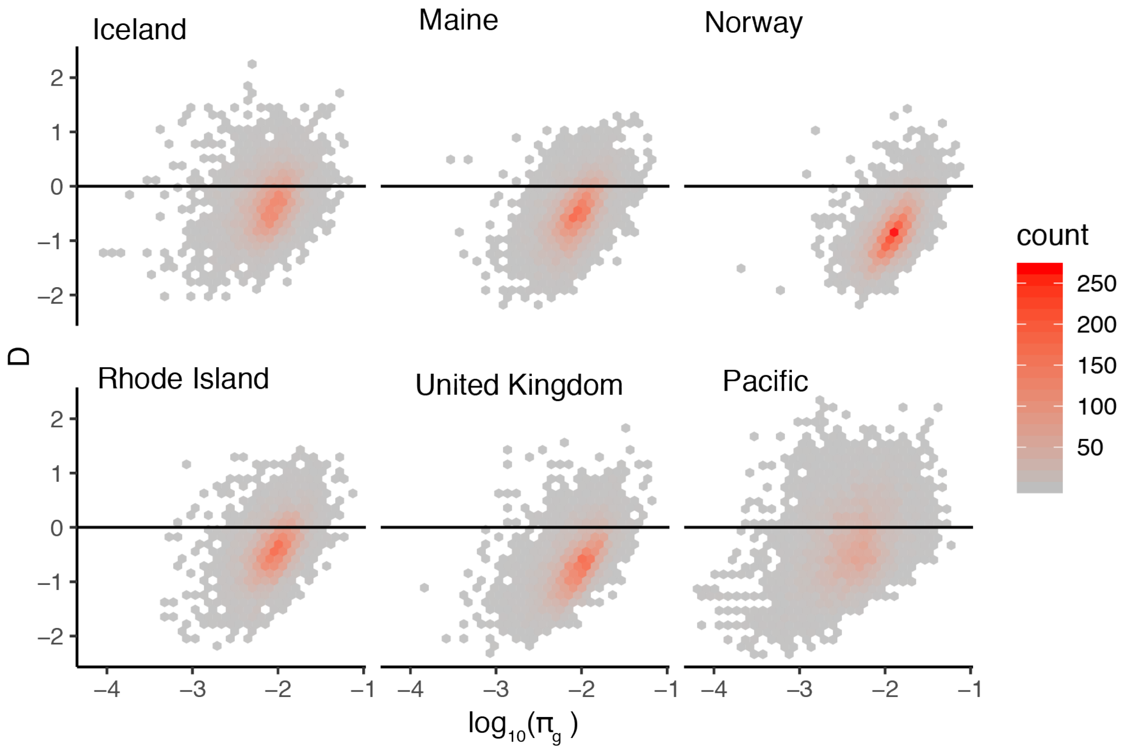

### Figure S3

A negative f3 statistic is evidence that the target population is admixed between source 1 and source 2 populations. Unsurprisingly, the outgroup WCAN was not found to be admixed between any other two populations. Remarkably, nearly all other f3 statistics were found to be significantly negative. This finding suggests widespread genetic exchange across the North Atlantic. Ranking the North American target populations by their f3 statistics, we found that UK and NOR had the most evidence of admixture, followed by ICE and RI, and finally by ME.

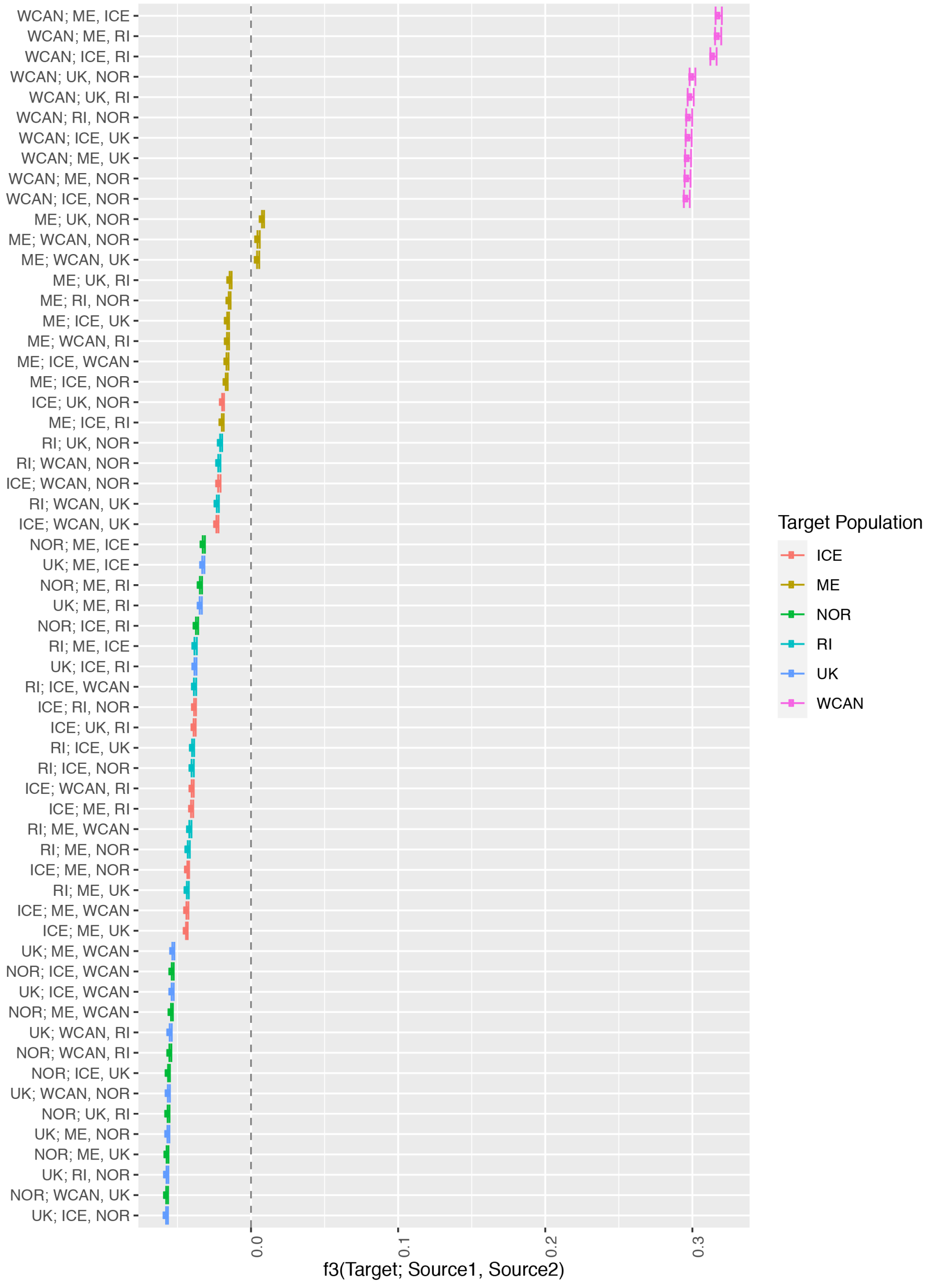

### Figure S4

Various plots of $F_{ST}$ across populations. A) $F_{ST}$ pairwise comparison for pool-seq populations. B) $F_{ST}$ tree of pool-seq populations. C) $F_{ST}$ distributions among pool-seq libraries. D) Example of a windowed $F_{ST\ plot}$ for comparisons within the Atlantic (blue lines) and between oceans (orange lines) of one of the longest scaffolds in Sbal3.1 (Scaffold #29).

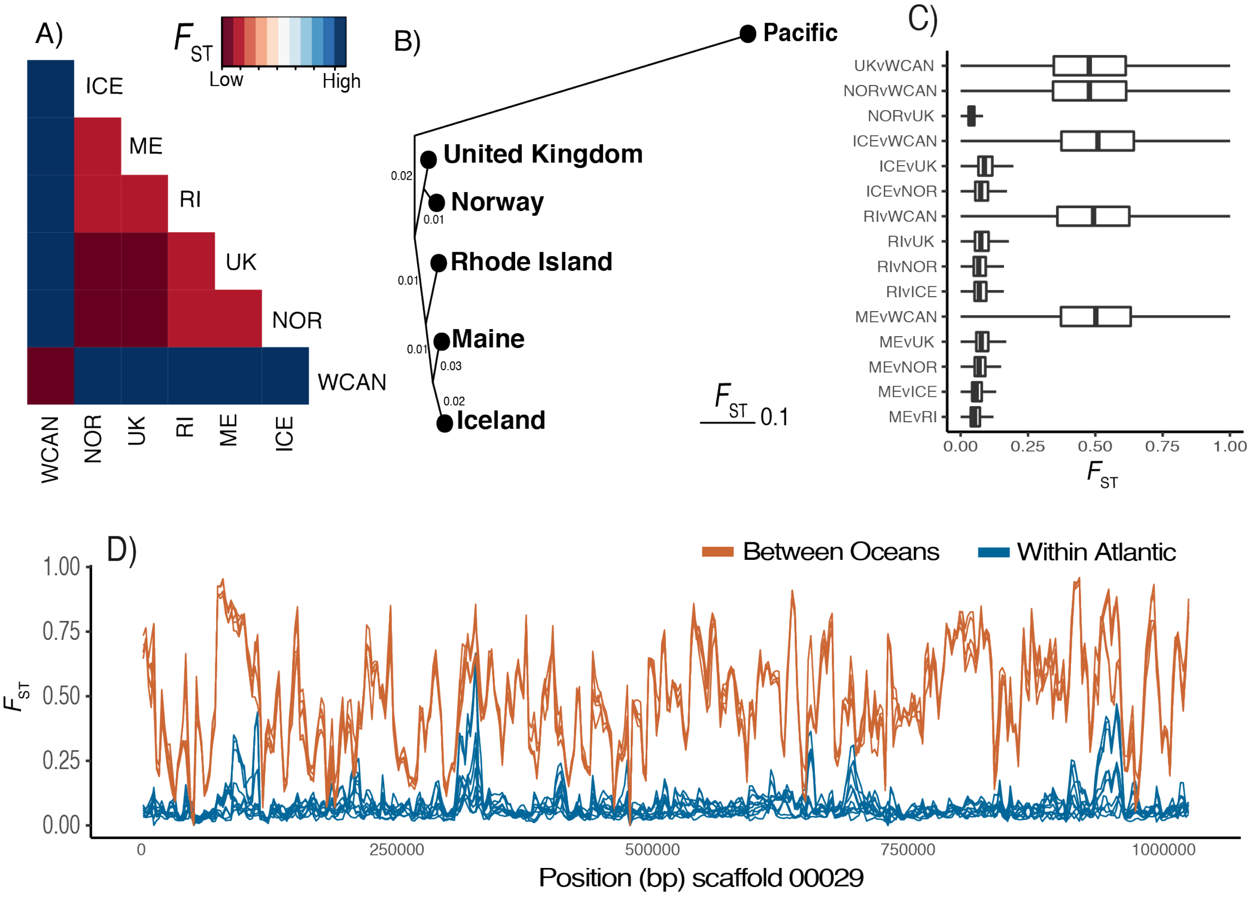

### Figure S5

Distribution of TSP SNP heterozygosity values across all populations. The grey box indicates rare variation.

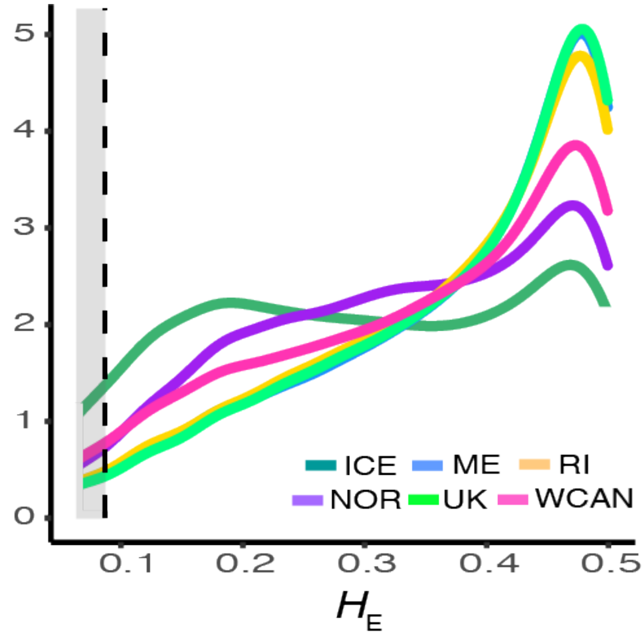

### Figure S6

Distribution of $\pi$ (A) and D (B) across populations as a function of exon length. Red lines indicate exons without a TSP. Green lines are regions with TSPs.

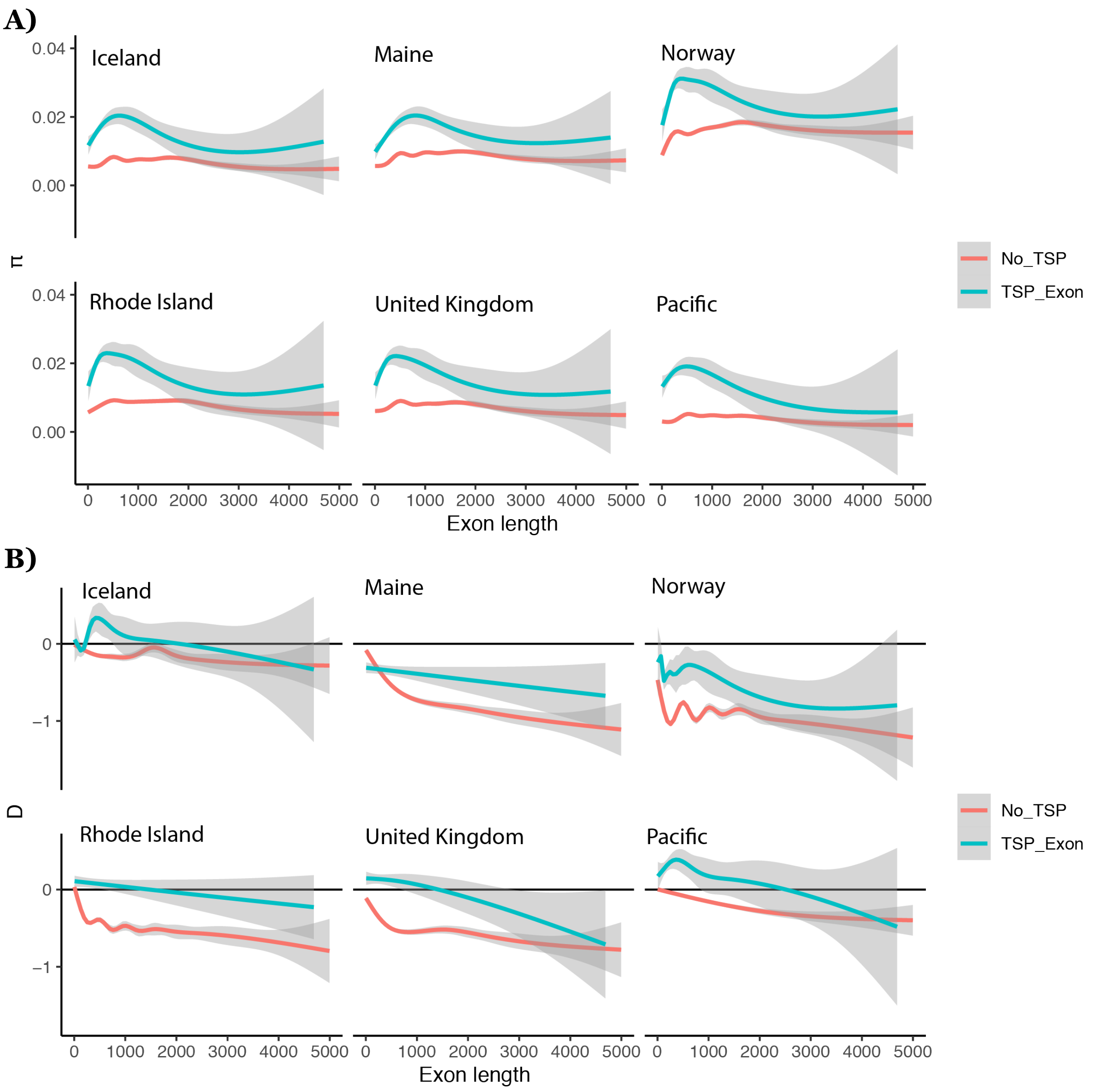

### Figure S7

Distribution of $F_{ST}$ values across populations as a function for exons with and without a TSP.

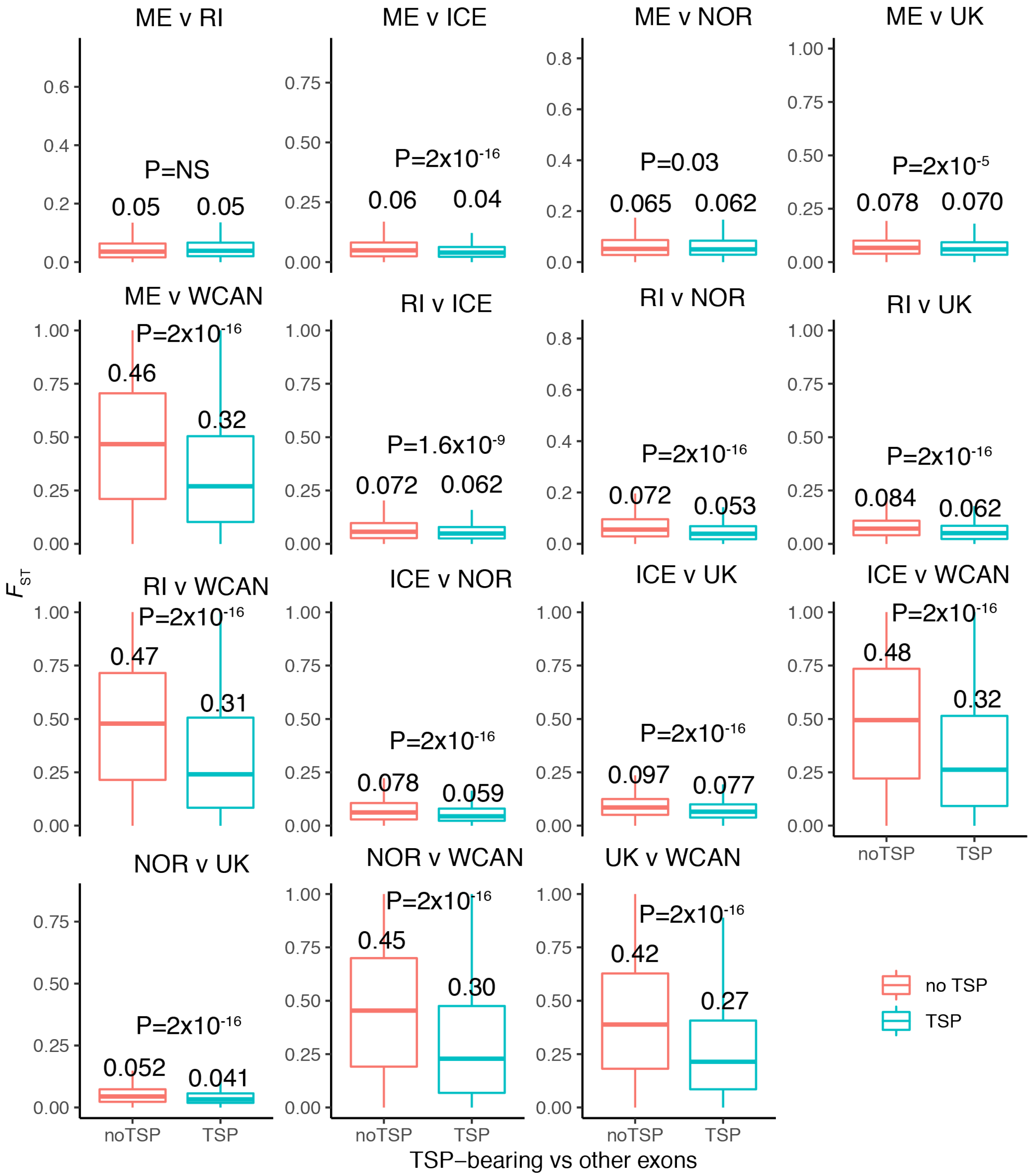

### Table S1

Various population genetic parameters across populations for the mtDNA COX I dataset. N: number of sequences. π: nucleotide diversity, $\pi_{se}$: standard error of nucleotide diversity, hd: haplotype diversity, $hd_{se}$ standard error of haplotype diversity, $R^2$: Ramos-Onsins-Rozas Test of Neutrality, FL-D: Fu and Li' D, FL-D(P): p-value of FL-D. The top table captures patterns in populations. The bottom table captures patterns in haplotypes.

| mtDNA Pop | N | $\pi$ | $\pi_{se}$ | hd | $hd_{se}$ | $R^2$ | FL-D | FL-D(P) |
|---|---|---|---|---|---|---|---|---|
| Maine | 21 | 0.007 | 0.001 | 0.81 | 0.08 | 0.115 | -0.175 | NS |
| Canada East | 13 | 0.008 | 0.00087 | 0.731 | 0.096 | 0.186 | 0.479 | NS |
| Rhode Island | 210 | 0.00872 | 0 | 0.83 | 0.016 | 0.043 | -3.705 | 0.02 |
| France | 7 | 0.009 | 0.001 | 1 | 0.076 | 0.105 | -0.081 | NS |
| Iceland | 28 | 0.007 | 0.001 | 0.627 | 0.084 | 0.136 | -0.252 | NS |
| Norway (Bergen) | 243 | 0.008 | 0 | 0.963 | 0.007 | 0.02 | -3.644 | 0.02 |
| Pacific | 25 | 0.00171 | 0.00039 | 0.687 | 0.101 | 0.052 | -2.71 | 0.05 |
| Wales | 27 | 0.005 | 0.001 | 0.775 | 0.086 | 0.06 | -1.836 | NS |
| Faroe | 20 | 0.003 | 0.001 | 0.795 | 0.087 | 0.105 | -0.858 | NS |
| Sweden | 114 | 0.008 | 0 | 0.959 | 0.012 | 0.029 | -2.996 | 0.02 |
| Ireland | 4 | 0.00502 | 0.00183 | 0.833 | 0.222 | 0.2764 | -1.50675 | NS |

| mtDNA Hap | N | $\pi$ | $\pi_{se}$ | hd | $hd_{se}$ | $R^2$ | D | P(D) |
|---|---|---|---|---|---|---|---|---|
| a | 101 | 0.00137 | 0.0003 | 0.434 | 0.063 | 0.0203 | -2.5 | $P < 0.001$ |
| b | 239 | 0.00322 | 0.00022 | 0.802 | 0.031 | 0.0196 | -2.47 | $P < 0.01$ |
| bRI | 48 | 0.00179 | 0.00791 | 0.473 | 0.091 | 0.0343 | -2.49 | $P < 0.01$ |
| bSC | 165 | 0.00417 | 0.00022 | 0.887 | 0.02 | 0.0165 | -246 | $P < 0.01$ |
| c | 25 | 0.00171 | 0.00039 | 0.687 | 0.101 | 0.052 | -2.17 | $P < 0.01$ |

### Table S2

Table of $f_3$ statistics for populations.

| $admix_{pop}$ | $source_{pop1}$ | $source_{pop2}$ | $f_3$ | $f_{3sd}$ | $z_{score}$ |
|---|---|---|---|---|---|
| UK | ICE | NOR | -0.0571052 | 0.000631084 | -90.4876 |
| NOR | WCAN | UK | -0.0570665 | 0.00063273 | -90.1909 |
| UK | RI | NOR | -0.0568383 | 0.000633807 | -89.6777 |
| NOR | ME | UK | -0.0567307 | 0.000637825 | -88.9439 |
| UK | ME | NOR | -0.056157 | 0.000637971 | -88.0244 |
| NOR | UK | RI | -0.0560494 | 0.000659389 | -85.0021 |
| UK | WCAN | NOR | -0.0558212 | 0.000697829 | -79.9927 |
| NOR | ICE | UK | -0.0557825 | 0.000639627 | -87.211 |
| NOR | WCAN | RI | -0.0548245 | 0.00064953 | -84.4064 |
| UK | WCAN | RI | -0.0545963 | 0.000680949 | -80.1768 |
| NOR | ME | WCAN | -0.0537377 | 0.000632171 | -85.005 |
| UK | ICE | WCAN | -0.0532564 | 0.00069003 | -77.1798 |
| NOR | ICE | WCAN | -0.0532177 | 0.000657947 | -80.8844 |
| UK | ME | WCAN | -0.0528283 | 0.000684 | -77.2343 |
| ICE | ME | UK | -0.0436477 | 0.000582608 | -74.9178 |
| ICE | ME | WCAN | -0.0432196 | 0.000669592 | -64.5461 |
| RI | ME | UK | -0.0430016 | 0.000667617 | -64.4106 |
| ICE | ME | NOR | -0.0426995 | 0.00060548 | -70.5218 |
| RI | ME | NOR | -0.0423203 | 0.000703932 | -60.1199 |
| RI | ME | WCAN | -0.0412335 | 0.000719139 | -57.3373 |
| ICE | ME | RI | -0.0400345 | 0.000713144 | -56.1381 |
| ICE | WCAN | RI | -0.0396558 | 0.000713775 | -55.5578 |
| RI | ICE | NOR | -0.0396553 | 0.00074144 | -53.4842 |
| RI | ICE | UK | -0.0393884 | 0.000701639 | -56.1376 |
| ICE | UK | RI | -0.0383159 | 0.00070852 | -54.0788 |
| ICE | RI | NOR | -0.0380489 | 0.000663801 | -57.3198 |
| RI | ICE | WCAN | -0.0380485 | 0.000761202 | -49.9847 |
| UK | ICE | RI | -0.0377699 | 0.0006941 | -54.4157 |
| RI | ME | ICE | -0.0376697 | 0.000787393 | -47.8411 |
| NOR | ICE | RI | -0.0367141 | 0.000734247 | -50.0024 |
| UK | ME | RI | -0.0341567 | 0.000748696 | -45.6217 |
| NOR | ME | RI | -0.0340491 | 0.000740296 | -45.994 |
| UK | ME | ICE | -0.0324381 | 0.000739657 | -43.8556 |
| NOR | ME | ICE | -0.0320636 | 0.000723329 | -44.3278 |

| | | | | | |
|---|---|---|---|---|---|
| ICE | WCAN | UK | -0.0228294 | 0.000711613 | -32.0812 |
| RI | WCAN | UK | -0.022562 | 0.000731437 | -30.8461 |
| ICE | WCAN | NOR | -0.0215454 | 0.00078804 | -27.3405 |
| RI | WCAN | NOR | -0.0215449 | 0.00074135 | -29.0618 |
| RI | UK | NOR | -0.02032 | 0.00068411 | -29.7028 |
| ME | ICE | RI | -0.0191921 | 0.000648576 | -29.5912 |
| ICE | UK | NOR | -0.0189806 | 0.00065177 | -29.1216 |
| ME | ICE | NOR | -0.0165271 | 0.000667536 | -24.7584 |
| ME | ICE | WCAN | -0.0160071 | 0.000705874 | -22.6769 |
| ME | WCAN | RI | -0.0156283 | 0.000704957 | -22.1692 |
| ME | ICE | UK | -0.0155789 | 0.000655164 | -23.7786 |
| ME | RI | NOR | -0.0145416 | 0.000655737 | -22.1759 |
| ME | UK | RI | -0.0138603 | 0.000676008 | -20.5031 |
| ME | WCAN | UK | 0.00481126 | 0.000740763 | 6.495 |
| ME | WCAN | NOR | 0.00514703 | 0.000778182 | 6.61417 |
| ME | UK | NOR | 0.00813999 | 0.000688629 | 11.8206 |
| WCAN | ICE | NOR | 0.296239 | 0.00199847 | 148.233 |
| WCAN | ME | NOR | 0.296759 | 0.00200263 | 148.185 |
| WCAN | ME | UK | 0.297095 | 0.00202799 | 146.498 |
| WCAN | ICE | UK | 0.297523 | 0.00202869 | 146.657 |
| WCAN | RI | NOR | 0.297846 | 0.00200646 | 148.443 |
| WCAN | UK | RI | 0.298863 | 0.00203014 | 147.213 |
| WCAN | UK | NOR | 0.300088 | 0.00193788 | 154.854 |
| WCAN | ICE | RI | 0.314349 | 0.00212305 | 148.065 |
| WCAN | ME | RI | 0.317535 | 0.00207463 | 153.056 |
| WCAN | ME | ICE | 0.317913 | 0.00214256 | 148.38 |

### Table S3

GO enrichment analysis of genes with positive $CPD_{w-b}$

##### Functions Ontology

| GO term | Description | P-value | Genes_ |
|---|---|---|---|
| GO:0004888 | transmembrane signaling receptor activity | 8.96E-6 | CG11318 - cg11318 gene product from transcript cg11318-ra<br>mthl10 - methuselah-like 10<br>Eph - eph receptor tyrosine kinase<br>Ir93a - ionotropic receptor 93a<br>KaiRIA - cg18039 gene product from transcript cg18039-ra<br>Ir25a - ionotropic receptor 25a<br>AlCR2 - allatostatin c receptor 2<br>Lar - leukocyte-antigen-related-like<br>ptc - patched<br>mGluRA - metabotropic glutamate receptor<br>plexA - plexin a<br>nAcRalpha-30D - nicotinic acetylcholine receptor alpha 30d |
| GO:0038023 | signaling receptor activity | 8.92E-5 | CG11318 - cg11318 gene product from transcript cg11318-ra<br>mthl10 - methuselah-like 10<br>Eph - eph receptor tyrosine kinase<br>Ir93a - ionotropic receptor 93a<br>KaiRIA - cg18039 gene product from transcript cg18039-ra<br>Ir25a - ionotropic receptor 25a<br>AlCR2 - allatostatin c receptor 2<br>ptc - patched<br>Lar - leukocyte-antigen-related-like<br>mGluRA - metabotropic glutamate receptor<br>plexA - plexin a<br>nAcRalpha-30D - nicotinic acetylcholine receptor alpha 30d |
| GO:0060089 | molecular transducer activity | 1.95E-4 | CG11318 - cg11318 gene product from transcript cg11318-ra<br>mthl10 - methuselah-like 10<br>Eph - eph receptor tyrosine kinase<br>Ir93a - ionotropic receptor 93a<br>KaiRIA - cg18039 gene product from transcript cg18039-ra<br>Ir25a - ionotropic receptor 25a<br>AlCR2 - allatostatin c receptor 2<br>ptc - patched<br>Lar - leukocyte-antigen-related-like<br>mGluRA - metabotropic glutamate receptor<br>plexA - plexin a<br>nAcRalpha-30D - nicotinic acetylcholine receptor alpha 30d |
| GO:0008066 | glutamate receptor activity | 7.96E-4 | Ir93a - ionotropic receptor 93a<br>Ir25a - ionotropic receptor 25a<br>KaiRIA - cg18039 gene product from transcript cg18039-ra<br>mGluRA - metabotropic glutamate receptor |
| GO:0022838 | substrate-specific channel activity | 9.84E-4 | wtrw - water witch<br>Ir93a - ionotropic receptor 93a<br>KaiRIA - cg18039 gene product from transcript cg18039-ra<br>Ir25a - ionotropic receptor 25a<br>pyx - pyrexia<br>pain - painless<br>nAcRalpha-30D - nicotinic acetylcholine receptor alpha 30d<br>Shaw - shaker cognate w |

| | | | |
|---|---|---|---|
| GO:0005216 | ion channel activity | 9.84E-4 | wtrw - water witch<br>Ir93a - ionotropic receptor 93a<br>KaiRIA - cg18039 gene product from transcript cg18039-ra<br>Ir25a - ionotropic receptor 25a<br>pyx - pyrexia<br>pain - painless<br>nAcRalpha-30D - nicotinic acetylcholine receptor alpha 30d<br>Shaw - shaker cognate w |

**Parts Ontology**

| GO term | Description | P-value | Genes |
|---|---|---|---|
| GO:0120025 | plasma membrane bounded cell projection | 8.14E-4 | Eph - eph receptor tyrosine kinase<br>Ir25a - ionotropic receptor 25a<br>AlCR2 - allatostatin c receptor 2<br>nAcRalpha-30D - nicotinic acetylcholine receptor alpha 30d<br>CG42629 - cg42629 gene product from transcript cg42629-rb<br>Shaw - shaker cognate w<br>Kif3C - cg17461 gene product from transcript cg17461-ra<br>shot - short stop<br>Lar - leukocyte-antigen-related-like<br>ptc - patched<br>CG8331 - cg8331 gene product from transcript cg8331-rd<br>Che-13 - hippi<br>arm - armadillo |
| GO:0042995 | cell projection | 9.08E-4 | Eph - eph receptor tyrosine kinase<br>Ir25a - ionotropic receptor 25a<br>AlCR2 - allatostatin c receptor 2<br>nAcRalpha-30D - nicotinic acetylcholine receptor alpha 30d<br>CG42629 - cg42629 gene product from transcript cg42629-rb<br>Shaw - shaker cognate w<br>Kif3C - cg17461 gene product from transcript cg17461-ra<br>shot - short stop<br>Lar - leukocyte-antigen-related-like<br>ptc - patched<br>CG8331 - cg8331 gene product from transcript cg8331-rd<br>Che-13 - hippi<br>arm - armadillo |
| GO:0034702 | ion channel complex | 9.32E-4 | wtrw - water witch<br>KaiRIA - cg18039 gene product from transcript cg18039-ra<br>pain - painless<br>pyx - pyrexia<br>nAcRalpha-30D - nicotinic acetylcholine receptor alpha 30d<br>Shaw - shaker cognate w |

### Table S4

GO enrichment analysis of genes with negative $CPD_{w-b}$

#### Processes Ontology

| GO term | Description | P-value | Genes |
|---|---|---|---|
| GO:0048646 | anatomical structure formation involved in morphogenesis | 3.78E-5 | S - star<br>hh - hedgehog<br>ds - dachsous<br>sls - sallimus<br>bt - bent<br>Nc - nedd2-like caspase<br>Trh - tryptophan hydroxylase<br>heph - hephaestus<br>lea - leak<br>exd - extradenticle<br>Pax - paxillin<br>kkv - krotzkopf verkehrt<br>Csk - c-terminal src kinase<br>mmy - mummy<br>Mhc - myosin heavy chain<br>unc-5 - cg8166 gene product from transcript cg8166-ra |
| GO:0032501 | multicellular organismal process | 9.8E-4 | Dhc36C - dynein heavy chain at 36c<br>Hexo2 - hexosaminidase 2<br>hh - hedgehog<br>Cirl - cg8639 gene product from transcript cg8639-rg<br>ds - dachsous<br>Ppt1 - palmitoyl-protein thioesterase 1<br>Nc - nedd2-like caspase<br>Trh - tryptophan hydroxylase<br>eag - ether a go-go<br>exd - extradenticle<br>GluClalpha - cg7535 gene product from transcript cg7535-rk<br>Pkd2 - polycystic kidney disease gene-2<br>osa - cg7467 gene product from transcript cg7467-re<br>Snr1 - snf5-related 1<br>grim - cg4345 gene product from transcript cg4345-ra<br>Eaat1 - excitatory amino acid transporter 1<br>mats - mob as tumor suppressor<br>S - star<br>ftz-f1 - ftz transcription factor 1<br>Tmhs - tetraspan membrane protein in hair cell stereocilia ortholog<br>sls - sallimus<br>bt - bent<br>ovo - cg6824 gene product from transcript cg6824-re<br>nonA - no on or off transient a<br>Cht6 - cg43374 gene product from transcript cg43374-rc<br>dpr6 - cg14162 gene product from transcript cg14162-re<br>lea - leak<br>Mhc - myosin heavy chain<br>Mad - mothers against dpp<br>dikar - cg42799 gene product from transcript cg42799-rg<br>ninaC - neither inactivation nor afterpotential c<br>MRP - multidrug-resistance like protein 1<br>SoxN - soxneuro<br>unc-5 - cg8166 gene product from transcript cg8166-ra |

**Functions Ontology**

| GO term | Description | P-value | Genes_ |
|---|---|---|---|
| GO:0042803 | protein homodimerization activity | 4.09E-4 | dbo - diablo<br>lea - leak<br>Mhc - myosin heavy chain<br>KLHL18 - cg3571 gene product from transcript cg3571-rc<br>CG8965 - cg8965 gene product from transcript cg8965-ra<br>ncd - non-claret disjunctional<br>grim - cg4345 gene product from transcript cg4345-ra<br>Nc - nedd2-like caspase |

**Parts Ontology**

| GO term | Description | P-value | Genes_ |
|---|---|---|---|
| GO:0005886 | plasma membrane | 1.14E-4 | mats - mob as tumor suppressor<br>S - star<br>Hexo2 - hexosaminidase 2<br>Ance-3 - cg17988 gene product from transcript cg17988-rb<br>hh - hedgehog<br>Tmhs - tetraspan membrane protein in hair cell stereocilia ortholog<br>Duox - dual oxidase<br>slif - slimfast<br>Nc - nedd2-like caspase<br>E23 - early gene at 23<br>CG3216 - cg3216 gene product from transcript cg3216-rd<br>CG31121 - cg31121 gene product from transcript cg31121-ra<br>eag - ether a go-go<br>Fur2 - furin 2<br>Pax - paxillin<br>beta-Spec - beta spectrin<br>CG9981 - cg9981 gene product from transcript cg9981-rc<br>pyd - polychaetoid<br>Ptp99A - protein tyrosine phosphatase 99a<br>MRP - multidrug-resistance like protein 1<br>grim - cg4345 gene product from transcript cg4345-ra<br>CG5549 - cg5549 gene product from transcript cg5549-rb<br>Mcr - macroglobulin complement-related<br>Eaat1 - excitatory amino acid transporter 1 |

### **Appendix 1:** The Genome of *S. balanoides* (version 3.1).

This supplementary appendix described details on the genome assembly of the barnacle, Sbal3.1. This version was assembled using a hybrid approach which combined PacBio reads (SRR10011818) with short reads (SRR10011819). Two different individuals from Maine had to be used, one for each technology, due to minimum starting DNA requirements. The resulting assembly is a considerable improvement over previous versions of the genome (Nunez, et al. 2018a; Nunez, et al. 2018b). The assembly reconstructed 486 Mbp into 16,596 scaffolds. The N50 of the assembly 56,748 bp and the longest scaffold is 1.3 Mb. We used Jellyfish(Marcais and Kingsford 2011), a *k-mer* decomposition method, to understand how much of the genome was assembled in Sbal3.1 relative to expected total size. This was run on the Illumina Fasta files to generate a kmer histogram (k=21; **Fig. T1.1A and T1.1B**). Our results predicted a haploid genome size of 1.05 Gb for *Semibalanus*. GenomeScope (Vurture, et al. 2017) was run on the kmer hisogram to estimate genome size, percentage of repetitive sequence, and heterozygosity. GenomeScope predicted a haploid genome size of 1.05Gb (**Table T1.1**), with 417Mb (39.7%) of unique sequence, and 634Mb of repetitive sequence (60.3%). The predicted high proportion of repetitive elements highlights the inherent challenge of assembling barnacle genomes. We assessed genome completeness using the BUSCO method (Simao, et al. 2015). Of all expected metazoan genes in Sbal3.1, 57% are complete, 10% are fragmented, and 33% are missing. In addition, we used *ab initio* methods to predict and annotate the genome. Overall, we predicted 14,374 genes across all scaffolds, 7,862 of which were successfully annotated using homology to the *Drosophila melanogaster* genome (Genomics 2014; Hoskins, et al. 2015). GenomeScope predicted the heterozygocity in the genome to be 1.64%. We also estimated genome size using two additional methods to gain consensus. We ran findGSE(Sun, et al. 2018) on the histogram produced by Jellyfish the same genome size estimate to GenomeScope, 1.05Gb. We next use a different approach of aligning the Illumina reads to the genome and creating a histogram of the coverage using the 'samtools depth' command. The coverage was determined as the peak in the resulting histogram. Genome size was estimated as (N*(1-e)/C), where N is the total number of bases, e is the error rate (from GenomeScope) and C is the coverage. The Genome size estimate was: (50739026942*.9983) / 35 = 1.44Gb, which is very similar to the estimated genome size

**Table T1.1: Properties of Sbal3.1**

| **Property** | **min** | **max** |
|---|---|---|
| Heterozygosity | 1.643% | 1.648% |
| Genome Haploid Length | 1,050,181,468 bp | 1,050,817,812 bp |
| Genome Repeat Length | 633,509,918 bp | 633,893,785 bp |
| Genome Unique Length | 416,671,550 bp | 416,924,026 bp |
| Model Fit | 92.46% | 98.71% |
| Read Error Rate | 0.17% | 0.17% |

of the sister species, *S. cariosus* (1.37Gb). BUSCO(Simao, et al. 2015) scores for genome completion are shown in **Fig. T1.1C**.

**Figure T1.1:** A) GenomeScope analysis of Sbal3.1. B) Distribution of *k-mer* coverages. C) BUSCO assessment of genome completion for Sbal3.1.

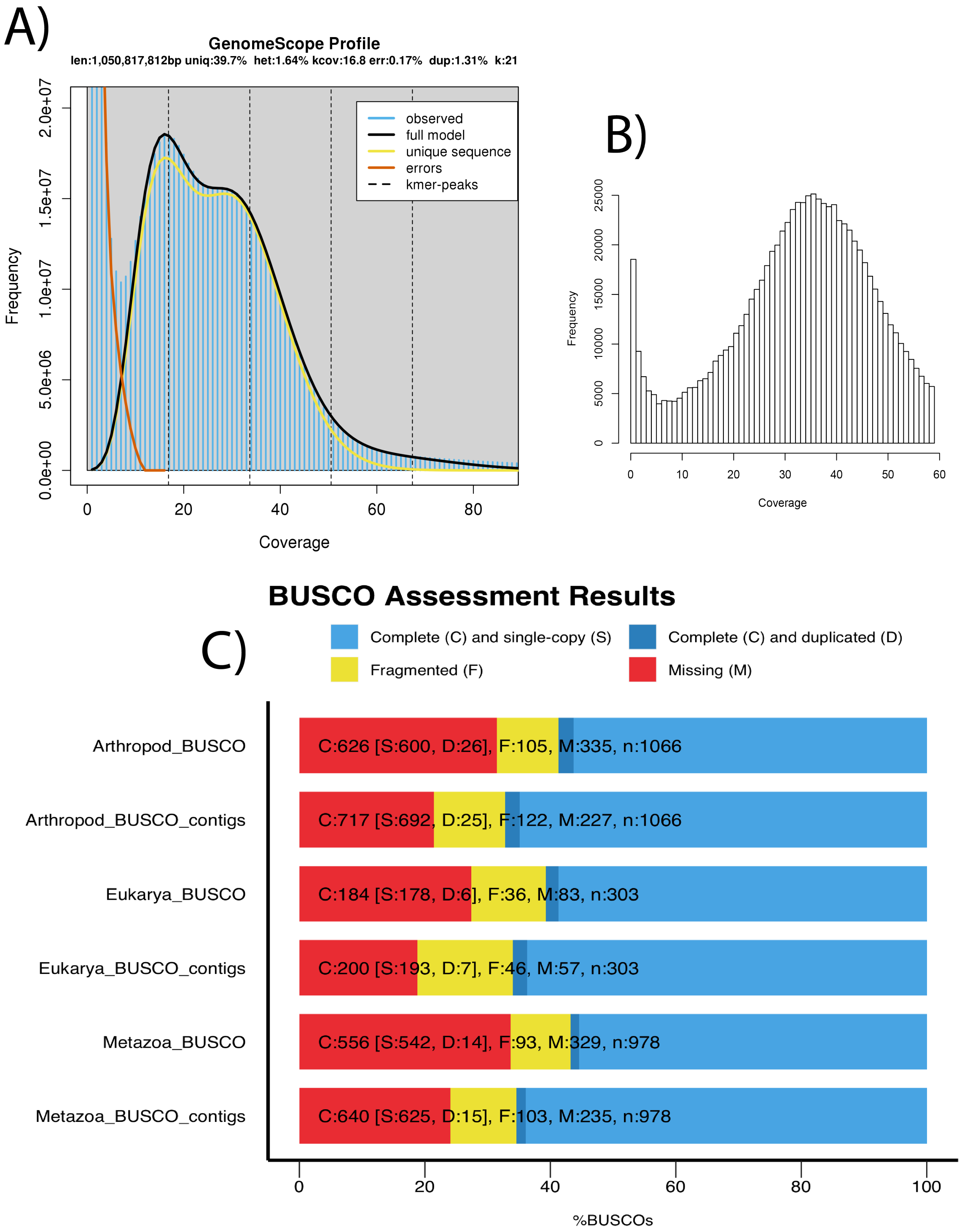

### **Appendix 2:** Details of standing genetic variation.

This appendix elaborated on details about the levels of standing variation observed in the species. First, we will describe the datasets used in the analysis (**Table T2.1**).

**Table T2.1:** various metrics of quality for all our genomic datasets

| Pop. | Type | SRA | Mapped Reads | Cov (mean) | Cov (sd) | IS (mean) | IS (sd) | GER | MQ |
|---|---|---|---|---|---|---|---|---|---|
| ICE | Pool | SRR10011808 | 190,417,336 | 48.5 | 364.4 | 281.9 | 331.1 | 0.0305 | 56.7 |
| ME | Pool | SRR10011798 | 207,966,897 | 54.4 | 382.0 | 354.7 | 321.4 | 0.0282 | 57.0 |
| RI | Pool | SRR10011813 | 284,914,248 | 72.5 | 657.1 | 182.3 | 390.3 | 0.0323 | 56.5 |
| UK | Pool | SRR10011805 | 313,449,000 | 81.3 | 483.0 | 198.6 | 356.9 | 0.0332 | 56.4 |
| NOR | Pool | SRR10011810 | 270,295,360 | 67.3 | 884.2 | 186.4 | 352.5 | 0.0344 | 56.2 |
| WCAN | Pool | SRR10011825 | 240,739,431 | 62.0 | 321.6 | 228.7 | 295.3 | 0.039 | 56.3 |
| ICE | Individual | SRR10011807 | 172,904,969 | 43.9 | 304.0 | 289.9 | 316.6 | 0.0299 | 56.7 |
| ME | Individual | SRR10011819 | 122,985,537 | 32.0 | 323.2 | 546.5 | 314.3 | 0.0225 | 57.4 |
| RI | Individual | SRR10011812 | 164,841,551 | 41.7 | 272.7 | 272.4 | 332.0 | 0.0301 | 56.7 |
| UK | Individual | SRR10011804 | 298,862,117 | 76.2 | 434.4 | 197.1 | 338.1 | 0.0329 | 56.3 |
| NOR | Individual | SRR10011809 | 265,634,252 | 67.5 | 575.3 | 192.0 | 338.3 | 0.0332 | 56.3 |
| WCAN | Individual | SRR10011814 | 244,314,278 | 63.3 | 323.3 | 240.1 | 294.9 | 0.0391 | 56.2 |

After normalizing the coverage of all libraries to 30X, we retained 3,798,898 high quality genome-wide SNPs observed either as polymorphisms or fixed differences within all populations and across species. Most SNPs are non-coding (83.3%) and 16.7% are coding. In term of non-coding loci, 57.4% are introns, 20.4% are intergenic, 3.3% are promoters, 1.10% are 5'UTR, and 0.84% are 3'UTR. For functional SNPs, relative to the whole dataset, 9.1% are synonymous, 6.7% are non-synonymous, 0.25% are non-translated, and 0.11% are non-sense.

We used all Pool-seq libraries to characterize genetic variation in the species across ocean basins. These libraries were mapped to our newly assembled *S. balanoides* genome. We estimated allele frequencies across single nucleotide polymorphisms (SNPs) genome-wide, as well as two metrics of genetic variation across the range of the species (**Table T2.2**).

**Table T2.2:** various metrics of standing variation in Sbal3.1. ***Npool:*** Number of individuals sequenced in the pool. ***S:*** Number of segregating sites segregating at common frequencies (> 5%). $S_{NS}$**:** Number of nonsynonymous common segregating sites. $S_S$**:** Number of synonymous common segregating sites. $\theta$**:** Mutation scaled effective population size (4Neμ). $\pi_g$**:** Nucleotide diversity at gene loci. $\pi_i$**:** Nucleotide diversity at intron loci. $\pi_e$**:** Nucleotide diversity at exon loci. $D_g$**:** Tajima's D at whole genes. $D_i$**:** Tajima's D in intron loci. $D_e$**:** Tajima's D in exon loci.

| Pop. | *Npool* | *S* | $S_{NS}$ | $S_S$ | $\theta$ | $\pi_g$ | $\pi_i$ | $\pi_e$ | $D_g$ | $D_i$ | $D_e$ |
|---|---|---|---|---|---|---|---|---|---|---|---|
| ICE | 20 | 635,268 | 41,447 | 60,965 | 0.99% | 0.92% | 1.01% | 0.73% | -0.31 | -0.27 | -0.14 |
| ME | 37 | 746,867 | 58,685 | 90,096 | 1.09% | 0.97% | 1.02% | 0.81% | -0.49 | -0.43 | -0.47 |

| | | | | | | | | | | | |
|---|---|---|---|---|---|---|---|---|---|---|---|
| RI | 38 | 686,354 | 47,968 | 75,413 | 1.11% | 1.01% | 1.06% | 0.85% | -0.39 | -0.30 | -0.36 |
| UK | 28 | 927,213 | 98,679 | 164,833 | 1.19% | 1.02% | 1.07% | 0.81% | -0.65 | -0.68 | -0.42 |
| NOR | 20 | 686,867 | 45,039 | 69,782 | 1.66% | 1.33% | 1.20% | 1.49% | -0.85 | -0.80 | -0.86 |
| WCAN | 20 | 305,547 | 24,383 | 29,141 | 0.58% | 0.55% | 0.60% | 0.50% | -0.29 | -0.16 | -0.09 |

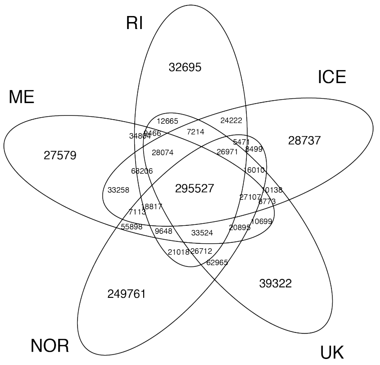

*Figure T2.1: Private variation in the Atlantic. Venn visualization*

We also estimated the number of shared and private mutations in the North Atlantic (**Fig. T2.1**): of all SNPs, 9.0% are present across the entire basin. ME, RI and ICE populations harbor the lowest number of private alleles (4.6-5.9%), and the UK population harbors 7.2% private SNPs. NOR, on the other hand, harbors an order of magnitude more private SNPs than any other Atlantic population (32.4%). In WCAN, 68% of all observed mutations are private to the basin. The number of shared alleles across oceans varies from 13% to 17%, with the UK sharing the most alleles (enrichment $P < 2.2\text{x}10^{-16}$).

We sought to classify what genes have consistent high and low levels of variation in the species. To this end, we applied a principal component analysis (PCA) using populations as variables and scaled values of $\pi_g$ as observations. The resulting PCA has a first dimension which robustly predicts the species wide levels of generic variation across all populations (**Fig. T2.2**; Pearson corr. = 0.872; $P < 2.2\text{x}10^{-16}$). We used the PCA projection quantiles between < 10$^{th}$, 10$^{th}$-20$^{th}$, 80$^{th}$-90$^{th}$, and > 90$^{th}$ to classify individual genes as having '*very low*', '*low*', '*high*', and '*very high*' levels of genetic variation. We then used gene ontology (GO) enrichment analysis on each classification. To avoid ascertainment bias, we tested enrichment relative to all annotated genes in Sbal3.1, and not against all annotated genes of Drosophila. For the $\pi_{very\ low}$, $\pi_{low}$, $\pi_{high}$ sets, we did not detect any significant, FDR corrected, GO enrichment. However, the $\pi_{very\ high}$ set shows statistically significant enrichment for the GO terms "*immune response*" (Process ontology, GO:0006955; FDR $P_{FDR} < 0.06$), and "*peptidase activity*" (Function

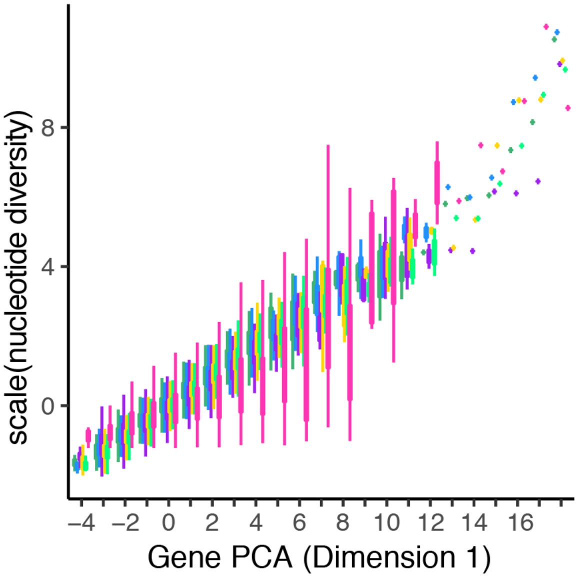

*Figure T2.2: Genetic variation scaled and sorted plotted against PC1 of the gene PCA. The colors are the same as in the main text. WCAN (pink), ME (blue), RI (yellow), ICE (dark green), NOR (purple), UK (light green).*

ontology, GO:0008233 and GO:0070011; $P_{FDR} < 0.005$). We selected 12 annotated genes within the set (**T2.3A**) and conducted and extended blast search against the entire National Center for Biotechnology Information's (NCBI) *nr* dataset (**T2.3B**), as assessing the robustness of their predicted tridimensional (3D) protein structure (**see Table. T2.3C**). These results provide confidence that these '*high variation*' loci are likely real genes, and not the result of faulty annotation, or spurious read-mapping to repetitive or chimerically assembled regions. These genes are homologs of *Mbo*, *Duox* (2 putative paralogs), *Tep2*, *Hml*, *Zir* (from the immunity GO term), as well as *Nep1*, *CYLD*, *stl*, *Tequila*, and *Ide* (from the peptidase GO term). Out of these 11 genes, 9 have a direct hit to a corresponding homolog in the closely related species (*Amphibalanus amphitrite*). Similarly, 10 of these genes show high confidence protein folding predictions.

**Fig. T2.3A**. High variation genes. Sbal3 Annotation

| Gene ID (Sbal) | Annotation name | Dmel equivalent | FBGN | GO term |
|---|---|---|---|---|
| g10782 | *Mbo* | CG6819 | FBgn0026207 | Immunity |
| g13789, g8744 | *Duox* | CG3131 | FBgn0283531 | Immunity |
| g14125 | *Tep2* | CG7052 | FBgn0041182 | Immunity |
| g6580 | *Hml* | CG7002 | FBgn0029167 | Immunity |
| g283 | *Zir* | CG11376 | FBgn0031216 | Immunity |
| g2658 | *Nep1* | CG5905 | FBgn0029843 | peptidase |
| g4577 | *CYLD* | CG5603 | FBgn0032210 | peptidase |
| g9458 | *stl* | CG3622 | FBgn0086408 | peptidase |
| g1844 | *Tequila* | CG4821 | FBgn0023479 | peptidase |
| g12306 | *Ide* | CG5517 | FBgn0001247 | peptidase |

**Fig. T2.3B**. High variation genes. Blast against the entire *nr* database

| Gene ID (Sbal) | Blast hit (all nr) | Taxon | e-value | Cover | Similarity |
|---|---|---|---|---|---|
| g10782 | N/A | N/A | N/A | N/A | N/A |
| g13789, | Dual oxidase 2 | *Amphibalanus amphitrite* | $10^{-45}$ | 100% | 74% |
| g8744 | dual oxidase 2-like | *Ixodes scapularis* | $10^{-157}$ | 88% | 49% |
| g14125 | Alpha-2-macroglobulin-like | *Amphibalanus amphitrite* | 0.0 | 98% | 72% |
| g6580 | Mucin-2 | *Amphibalanus amphitrite* | $10^{-163}$ | 98% | 78% |
| g283 | Dedicator of cytokinesis protein 7 | *Amphibalanus amphitrite* | $10^{-163}$ | 76% | 59% |
| g2658 | Neprilysin-11 | *Amphibalanus amphitrite* | 0.0 | 99% | 53% |
| g4577 | Ubiquitin carboxyl-terminal hydrolase CYLD | *Amphibalanus amphitrite* | $10^{-39}$ | 100% | 94% |
| g9458 | A disintegrin and metalloproteinase with | *Amphibalanus amphitrite* | $10^{-84}$ | 100% | 94% |

| | | | | | |
|---|---|---|---|---|---|
| | thrombospondin motifs adt-2 | | | | |
| g1844 | Lysyl oxidase 2 | *Amphibalanus amphitrite* | $10^{-96}$ | 89% | 44% |
| g12306 | Insulin-degrading enzyme | *Amphibalanus amphitrite* | 0.0 | 77% | 68% |

**Fig. T2.3C**. High variation genes. 3D protein prediction identification

| **Gene ID (Sbal)** | **Confidence** | **% of sequence modeled** | **PDB template** |
|---|---|---|---|
| g10782 | 98% | 50% | c5ijnF_ |
| g13789 | 99% | 91% | c3dd4A_ |
| g8744 | 100% | 44% | c5ooxA_ |
| g14125 | 100% | 95% | c2pn5A_ |
| g6580 | 99.3% | 72% | c4xbmB_ |
| g283 | N/A | N/A | N/A |
| g2658 | 100% | 79% | d1dmta_ |
| g4577 | 98% | 98% | c3wxeA_ |
| g9458 | 100% | 99% | c6qigA_ |
| g1844 | 100% | 23% | c5ze3B_ |
| g12306 | 100% | 98% | c2jbuB_ |

### **Appendix 3:** Bayesian Clock Analysis

This appendix elaborates on details about our Bayesian clock analysis. Our DNA analysis shows that populations in the Pacific and Atlantic oceans are highly diverged (**Fig. 1D**). This divergence suggests that populations across basins have not exchanged migrants in a long time. Current models of historical phylogeography posit that *S. balanoides* first speciated in the North Pacific 8-13 mya(Perez-Losada, et al. 2008; Herrera, et al. 2015). The species then invaded the North Atlantic Ocean during the events of the trans-Arctic interchange 1-3 mya (Vermeij 1991). We revisited this classic phylogeography hypothesis using our *COX I* dataset. First, we characterized haplotypes in our data. We observed multiple haplogroups previously characterized in the North Atlantic (named "*a*" and "*b*"; (Brown, et al. 2001; Wares and Cunningham 2001; Flight, et al. 2012; Nunez, et al. 2018b)). Notably, the inclusion of North Pacific samples reveals the existence of a previously unobserved haplogroup "*c*" (**Fig. 1D-inset**). This haplogroup is only observed in the Pacific and is highly differentiated from any of its Atlantic counterparts ($F_{ST}$ ~ 0.6; **Fig. S4**). We used a Bayesian molecular clock analysis (22) on the *COX I* data to estimate divergence times between Pacific and Atlantic populations. For this analysis, we used the mutation rate calculated by Wares and Cunningham(Wares and Cunningham 2001) and a strict clock model. Additionally, we only considered 3$^{rd}$ codon positions and implemented the HKY site model with estimated frequencies as well as the calibrated Yule model. The prior for the divergence date was a uniform distribution from 0 to 3.5 mya. All other priors were left in their default settings. All estimates were run with a MCMC chain length of 100 million with a pre-burnin of 25 million and checked for convergence by visual inspection of the log files produced by BEAST with Tracer v1.6. We estimated a divergence time between oceans of 1.91 my (95% Highest Posterior Density [HPD] interval = 0.66-3.29 my). Given the broad interval of these estimates, we repeated the analysis using a separate dataset composed of all protein-coding genes from the complete mitochondrial genomes from *S. balanoides* (from Atlantic and Pacific individuals), *S. cariosus, Balanus balanus, Megabalanus ajax, Megabalanus volcano,* as well as an outgroup to barnacles, the copepod *Tigriopus californicus*. The sequences of *Semibalanus* from North Atlantic individuals, as well as *Balanus* and *Megabalanus* are available in NCBI (**see extended methods**). The sequences for the Pacific *S. balanoides* and *S. cariosus* individuals were assembled *de novo* for this analysis (**see extended methods).** As above, we implemented a strict clock model using the Wares and Cunningham mutation rate as well as a uniform prior from 0-3.5 mya and only considered 3$^{rd}$ positions across the whole molecule with a MCMC chain length of 100 million with a pre-burnin of 25 million. In this analysis, our estimate of Pacific-Atlantic divergence is 1.95 my (95% HPD = 1.58-2.32 my). If the *Tigriopus* outgroup is excluded, this number changes to

1.84 my (95% HPD = 1.47-2.22 my). Overall, both the COX I and the complete mtDNA analysis produce congruent estimates of trans-Arctic divergence.

### **Appendix 4:** ABBA/BABA

This appendix elaborated on details about our ABBA-BABA tests for gene tree heterogeneity. ABBA-BABA tests for gene tree heterogeneity were done for triplet taxa subsampled from the Atlantic *S. balanoides* population (**see Dataset S3**). The analysis revealed multiple instances of potential gene flow between non-sister populations. Most notably, the significantly positive D-statistic for the ((ICE, ME), RI) triplet indicates gene flow between ME and RI since the divergence between ME and ICE (**Table T4.1**, **Fig. T4.1**) In this case, the f4-ratio indicates an admixture proportion of approximately 2.7 % (**Table T4.1**.) and the lineage-specific f-branch statistics suggests that the donor population of this exchange is ME (**Fig. T4.2**).

**Table T4.1:** The D-statistic and $f_4$-ratio for all possible 3-taxon "triplet" rearrangements that can be subsampled from the inferred true phylogeny of five Atlantic *S. balanoides* populations, with the Canada Pacific population used to assign the ancestral state of the outgroup. For each triplet, P1 and P2 are the pair with the highest count of BBAA sites (i.e. sister taxa in the triplet). Bolded p-values are significant after accounting for multiple comparisons with the Benjamini-Hochberg procedure with false discovery rate of 0.05. Red: evidence for gene flow between RI and ME since the split of ICE and ME. Blue: evidence for gene flow between RI and (NOR+UK) that is independent of (ME+ICE), although note that other scenarios are possible.

| P1 | P2 | P3 | D | p-value | $f_G$ |
|---|---|---|---|---|---|
| NOR | UK | ICE | 0.0044 | 0.0407 | 0.0066 |
| NOR | UK | ME | 0.0051 | 0.0080 | 0.0069 |
| ME | ICE | NOR | 0.0023 | 0.1399 | 0.0034 |
| ICE | RI | NOR | 0.0066 | **0.0060** | 0.0103 |
| ME | RI | NOR | 0.0091 | **0.0000** | 0.0137 |
| ICE | ME | RI | 0.0150 | **0.0000** | 0.0271 |
| NOR | UK | RI | 0.0028 | 0.0858 | 0.0042 |
| ME | ICE | UK | 0.0016 | 0.2138 | 0.0023 |
| ICE | RI | UK | 0.0049 | **0.0254** | 0.0075 |
| ME | RI | UK | 0.0066 | **0.0007** | 0.0098 |

Also, evident in the ABBA-BABA results are four triplet sets with significantly positive D-statistics that may indicate as few as one ancestral gene flow event (**Table T4.1,** blue rows), albeit with a low proportion of introgressed material (0.7 % to 1.3 %) (**Table T4.1**., **Fig. T4.1**). While these tests are consistent with the hypothesis of four separate introgression events, shared ancestry among the taxa means that other scenarios are possible. Gene flow may have occurred between RI and NOR since the splits between RI+ME and RI+ICE; and similarly, between RI and UK since the splits between RI+ME and RI+ICE. Yet, these results may be reconciled by a single gene flow between the ancestor of UK+NOR and RI before the split of ICE +ME, seeing as both ICE+ME and UK+NOR are sister populations. To further complicate interpretation of these results, any ongoing gene flow between sister populations (e.g., NOR+UK) may muddle signals of gene flow that was, at one point, specific to either lineage of sister pairs. To make better sense of these correlated D-statistics and $f_4$-ratios, we considered lineage-specific f-branch statistics. These results indicate multiple instances of directional gene flow, including from RI to NOR (and a lesser extent to UK), and from UK to RI (and a lesser extent to ME and ICE) (**Fig. T4.2**). However we note that these correlated f-branch statistics, as with the D-statistics, may be products fewer actual introgression events(Malinsky, et al. 2020).

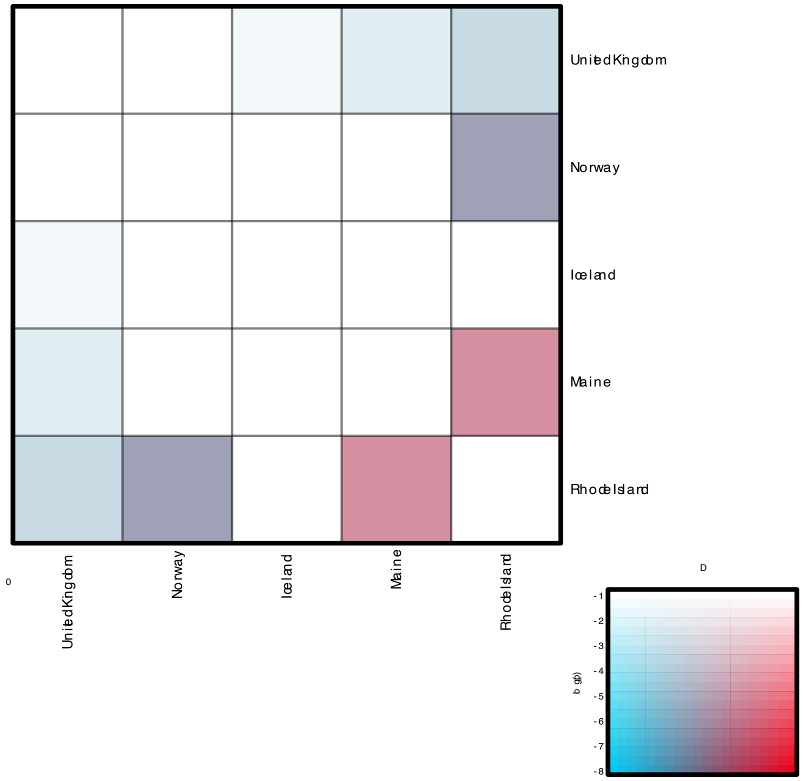

**Figure T4.1.** A graphical illustration of D-statistics corresponding to Table 1. Populations in positions P2 and P3 are sorted on the horizontal and vertical axes, and the color of the corresponding heatmap cell indicates the most significant D-statistic found with these two populations, across all possible populations in P1. More saturated colours indicate greater significance as shown in the inset (D and log(p)). Note that the highest D value shown on the inset is 0.02 and that the matrix is symmetric.

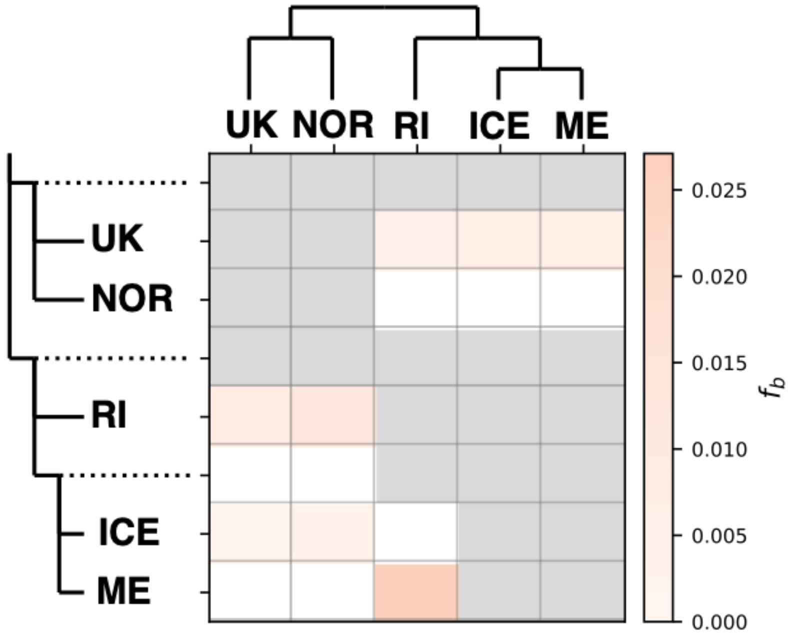

**Figure T4.2.** $F_{branch}$ results. The matrix shows the inferred f-branch statistics, showing excess allele sharing between the branch of the 'laddered' tree on the y axis (relative to its sister branch) and the population identified on the x axis.

### Supplementary Methods

**Sampling efforts.** This section indicates where barnacles were collected, as well as the GenBank accession Id. ME barnacles were collected in the Damariscotta River, near Hodgson's Island and the lower narrows south of the Darling Marine Center in Walpole, ME, USA. 37 individual were collected and sequenced as a pool (**SRR10011798**), with one sequenced to deep coverage (**SRR10011819**). RI samples were collected near the southern intersection of Conanicus Avenue and Bay View Drive in Jamestown, RI. 38 barnacles were collected and sequenced as a pool (**SRR10011813**), with one sequenced to deep coverage (**SRR10011812**). In Iceland, samples were collected on Sudurstrond at lat 64.155264 lon 22.022454 near Reykjavik. 20 individuals were sequenced as a pool (**SRR10011808**), and one in a single lane (**SRR10011807**). In the UK, samples were collected near the Porthcawl Sea Front in Wales. 34 individuals were sequenced as a pool (**SRR10011805**), and one in a single lane (**SRR10011804**). In the NOR, samples were collected near the Norddal, in *Møre* and *Romsdal* county. 20 individuals were sequenced as a pool (**SRR10011810**), and one in a single lane (**SRR10011809**). In the WCAN, samples were collected on Culvert Island, British Columbia. 20 individuals were sequenced as a pool (**SRR10011825**), and one in a single lane (**SRR10011814**). Species identity was confirmed using COX I sequencing. In addition, we sequenced COX I amplicons for 118 additional individuals from Tjärnö, Sweden; 245 from Norway, in Herdla Island, in Askøy near Bergen; and 20 from Tórshavn in the Faroe Islands. All samples can be accessed under GenBank ids: **MT329074-MT329592**.

**Genome assembly/ Mapping and filtering.** Scripts of the genome assembly and read mapping can be found, as used, in the github: **https://github.com/Jcbnunez/BarnacleEcoGenomics.** Note that reads mapping to multiple targets were removed using the XA tag. Overall, we scored sites, including sites in the sister taxa, *S. cariosus,* enforcing a genotyping call error < $1x10^{-10}$. enforcing a minimum coverage of 10X, a maf > 5%, and minimum quality score of 40. In all cases, we normalized the coverage of Pool-seq HTS libraries to 30X.

**Bayesian clock.** Both the Pacific *S. balanoides* and *S. cariosus* assemblies were performed with the same pipeline. Raw paired-end Illumina reads were filtered to remove low-complexity and low-quality reads. Initial assemblies were created using the bait and iterative method of MITObim v.1.9.1 (Hahn, et al. 2013). The Maine mtDNA assembly from Nunez *et al.* (Nunez, et al. 2018b) was used as a seed for the bait/iteration process (NCBI GenBank number: MG010647.1). The Pacific *S. balanoides* assembly took 5 iterations and the *S. cariosus* assembly took 30 iterations. The initial assemblies were annotated with MITOS, manually curated, and then annotated using Exonerate v2.2.0. Final assemblies were validated by constructing a maximum-likelihood phylogeny in iq-tree (Chernomor, et al. 2016) using the existing three assemblies of *S. balanoides* from Maine, Rhode Island, and Iceland (NCBI GenBank numbers: MG010647.1, MG010648.1, and MG010649.1), the complete mitochondrial sequences of *Balanus balanus* (NCBI GenBank number: KM660676.1), *Tigriopus californicus* (a copepod outgroup; NCBI GenBank number: NC_008831.2), *Megabalanus ajax* (NCBI GenBank number: NC_024636.1), and *Megabalanus volcano* (NCBI GenBank number: NC_0062931.2)

**Population genetic analyses.** We used the treepop program within TreeMix (version1.13) (Pickrell and Pritchard 2012) to estimate f3 (X; Y, Z) statistics, using blocks of 1000 SNPs (-k parameter) to account for LD, in order to explore evidence that the target population X is admixed between source populations Y and Z. To test for the occurrence of contemporary and historical gene flow between Atlantic *S. balanoides* populations, we performed ABBA-BABA tests and calculated D-statistics (Green *et al.* 2010) and related summary statistics using the program *Dsuite* (Malinsky, et al. 2020). We used biallelic SNPs obtained from whole genome DNA sequence data from one individual per sampled Atlantic *S. balanoides* populations. We performed tests of asymmetrical allele sharing between sister taxa in all possible 3-taxon "triplet" rearrangements that can be subsampled from the inferred true phylogeny: (((RI,(ME,ICE)),(UK,NOR)), Outgroup). For the outgroup (O), we used the sample from the Canada Pacific population of *S. balanoides*. Each triplet (((P1,P2),P3), O) was selected so that P1 and P2 share the derived allele (B) at more sites than do P1 and P3 or P1 and P2 – in other words, P1 and P2 are sister taxa in the triplet. With the ancestral state (A) determined by the outgroup, *D* is calculated as $(C_{ABBA} - C_{BABA}) / (C_{ABBA} + C_{BABA})$, where *C* refers to the count of sites conforming to a particular site pattern. Thus, in the absence of directional gene flow between P1 or P2 and P3, *D* is expected to be equal to zero. We present results for triplets with P1 and P2 ordered such that $C_{ABBA} \geq C_{BABA}$, so that D ≥ 0, for simplicity. Significantly positive D-scores show that derived alleles are not shared between the triplet as expected if they are related by a single species tree. Positive D-scores are thus interpreted as evidence of gene flow between P2 and P3 since the split between P1 and P2. Statistical significance of D is estimated using a standard block-jackknife procedure with sites partitioned into 20 bins. Z-scores were calculated as *D* over its standard error. We report the Z-score associated p-values and we accounted for multiple comparisons by applying the Benjamini-Hochberg procedure with a false discovery rate of 0.05. The $f_4$-ratio was calculated following the definition of (Patterson, et al. 2012) with the difference that P3 alleles were drawn at random from a diploid individual rather than a population of P3 individuals. The f4 ratio is expected to increase linearly with the proportion of the genome introgressed (Patterson, et al. 2012).

To gain further insight into the timing and directionality of historical gene flow, we further used the Dsuite package (Malinsky, et al. 2020) to investigate non-independent $f_4$-ratio results obtained from topology subsampling, which implements the f-branch statistic (Malinsky, et al. 2018). As described in the program manual, the f-branch statistic, $f_b(C)$, is a summary of *f* scores that captures excess allele sharing between a population C and descendants of the branch leading to population B (branch b) compared to the sister of branch *b*. This is calculated over all $f_4$-ratio results which had populations A and B as P1 or P2 and population C as P3, for the population trees that fit ((A,B),C). Thus, $f_b(C)$ scores are specific to a particular branch, b, denoted by the y-axis in Figure T4.2. MSMC analyses were run using parameters *–fixedRecombination,* -t 6, -p *10*1+15*2,* and a generation time of 1 year.

**Cophenetic distances.** The topological signatures of genes without a history of ancient balancing selection can be produced by multiple causes. In order to distinguish between genes with young haplotypes and genes with excess homozygotes, we carried out an additional topological contrast for each gene without evidence of ancient balancing selection. We distinguish genes with excess homozygosity as being those where the alleles from the Canadian population are not the most distantly related to all other alleles. We estimate this using the cophenetic branch length distances of the consensus trees (based on 100 bootstraps). The code for this is in **https://github.com/Jcbnunez/BarnacleEcoGenomics.**

### Dataset S1: Headers

Chr: Scaffold in Sbal3
Pos: position in scaffold
Rc: reference call
allele_count: number of alleles
allele_states: allele states
snp_type: variant type (relative to rc or in populations)
major_alleles.maa.: major allele in populations
MEHILC_ma: major allele in Maine
RIDLN_ma: major allele in Rhode Island
ICE_ma: major allele in Iceland
UKW_ma: major allele in UK
NOR_ma: major allele in Norway
WCAN_ma: major allele in Pacific Canada
CAR_ma: major allele in Cariosus
minor_alleles.mia.: minor allele in populations
N_Atl: Number of N calls
Pac: Call in the Pacific
Scar: Call in S. cariosus
MEHILC_mi: minor allele in Maine
RIDLN_mi: minor allele in Rhode Island
ICE_mi: minor allele in Iceland
UKW_mi: minor allele in UK
NOR_mi: minor allele in Norway
WCAN_mi: minor allele in Pacific Canada
CAR_mi: minor allele in Cariosus
MEHILC_count: count of the alternative allele in Maine
MEHILC_cov: coverage in Maine
RIDLN_count: count of the alternative allele in Rhode Island
RIDLN_cov: coverage in Rhode Island
ICE_count: count of the alternative allele in Iceland
ICE_cov: coverage in Iceland
UKW_count: count of the alternative allele in UK
UKW_cov: coverage in UK
NOR_count: count of the alternative allele in Norway
NOR_cov: coverage in Norway
WCAN_count: count of the alternative allele in Pacific Canada
WCAN_cov: coverage in Pacific Canada
CAR_count: count of the alternative allele in Cariosus
CAR_cov: coverage in Cariosus
QUAL: Quality of genotype call in Cariosus
Gt: Genotype in Cariosus
MissingDat_Natl: Missing data in the North Atlantic
LOCATION: Genomic element type
GENEID: Gene ID in Sbal3
SNP_id: SNP id
CONSEQUENCE: Synonymous/Nonsynonymous
<ME,RI,ICE...>_p_car: Allele frequency polarized to Cariosus
<ME,RI,ICE...>_p_WCA: Allele frequency polarized to Canada
<ME,RI,ICE...>_p_rc: Allele frequency polarized to Sbal3
<ME,RI,ICE...>_he: TSP SNP heterozygocity
Entry_gene_name: Annotation
PrimaryFBgn: FlyBase number
Type: TSP type

### Dataset S2: Headers

Gene: Gene ID in Sbal3
Mean_Feature_Posterior: Mean posterior for gene feature predictions
CPDwd: Mean Cophenetic distance (100 bootstraps)
CPDwd_stderr: Cophenetic distance standard error
CPDwd_min: Cophenetic distance minimum among all bootstraps
CPDwd_q25: distance $25^{th}$ quantile among all bootstraps
CPDwd_q75: distance $75^{th}$ quantile among all bootstraps
CPDwd_p_val: P-value
CPDwd_p_adjust: P-value adjusted (Bonferroni)
CPDwd_signif: Significance tag
Droso_gene_name: Annotation
PrimaryFBgn: FlyBase number

### Dataset S3: Headers

Chr: Scaffold in Sbal3
windowStart: start of the window
windowEnd: end of the window
D: ABBA/BABA statistic
f_d and f_dM: f statisitcs.

#### Supplementary References

Brown AF, Kann LM, Rand DM. 2001. Gene flow versus local adaptation in the northern acorn barnacle, Semibalanus balanoides: insights from mitochondrial DNA variation. Evolution 55:1972-1979.

Chernomor O, von Haeseler A, Minh BQ. 2016. Terrace Aware Data Structure for Phylogenomic Inference from Supermatrices. Syst Biol 65:997-1008.

Flight PA, O'Brien MA, Schmidt PS, Rand DM. 2012. Genetic Structure and the North American Postglacial Expansion of the Barnacle, Semibalanus balanoides. Journal of Heredity 103:153-165.

Genomics TFCBDGPC. 2014. Drosophila melanogaster (assembly Release 6 plus ISO1 MT). NCBI GenBank. https://www.ncbi.nlm.nih.gov/genome/47?genome_assembly_id=204923 Deposited August 2014.

Hahn C, Bachmann L, Chevreux B. 2013. Reconstructing mitochondrial genomes directly from genomic next-generation sequencing reads--a baiting and iterative mapping approach. Nucleic Acids Research 41:e129.

Herrera S, Watanabe H, Shank TM. 2015. Evolutionary and biogeographical patterns of barnacles from deep-sea hydrothermal vents. Mol Ecol 24:673-689.

Hoskins RA, Carlson JW, Wan KH, Park S, Mendez I, Galle SE, Booth BW, Pfeiffer BD, George RA, Svirskas R, et al. 2015. The Release 6 reference sequence of the Drosophila melanogaster genome. Genome Research 25:445-458.

Malinsky M, Matschiner M, Svardal H. 2020. Dsuite - fast D-statistics and related admixture evidence from VCF files. bioRxiv (unpublished data) https://www.biorxiv.org/content/10.1101/634477v2 - last accessed May 1, 2020.

Malinsky M, Svardal H, Tyers AM, Miska EA, Genner MJ, Turner GF, Durbin R. 2018. Whole-genome sequences of Malawi cichlids reveal multiple radiations interconnected by gene flow. Nat Ecol Evol 2:1940-1955.

Marcais G, Kingsford C. 2011. A fast, lock-free approach for efficient parallel counting of occurrences of k-mers. Bioinformatics 27:764-770.

Nunez JCB, Elyanow RG, Ferranti DA, Rand DM. 2018a. The Genome of Semibalanus balanoides, version 2.0. NCBI GenBank. https://www.ncbi.nlm.nih.gov/nuccore/PHFM00000000.1 Deposited Nov 2, 2018.

Nunez JCB, Elyanow RG, Ferranti DA, Rand DM. 2018b. Population Genomics and Biogeography of the Northern Acorn Barnacle (Semibalanus balanoides) Using Pooled Sequencing Approaches. In: Oleksiak MF, Rajora OP, editors. Population Genomics: Marine Organisms: Springer, Cham. p. 139-168.

Patterson N, Moorjani P, Luo Y, Mallick S, Rohland N, Zhan Y, Genschoreck T, Webster T, Reich D. 2012. Ancient admixture in human history. Genetics 192:1065-1093.

Perez-Losada M, Harp M, Hoeg JT, Achituv Y, Jones D, Watanabe H, Crandall KA. 2008. The tempo and mode of barnacle evolution. Mol Phylogenet Evol 46:328-346.

Pickrell JK, Pritchard JK. 2012. Inference of population splits and mixtures from genome-wide allele frequency data. PLoS Genet 8:e1002967.

Simao FA, Waterhouse RM, Ioannidis P, Kriventseva EV, Zdobnov EM. 2015. BUSCO: assessing genome assembly and annotation completeness with single-copy orthologs. Bioinformatics 31:3210-3212.

Sun H, Ding J, Piednoel M, Schneeberger K. 2018. findGSE: estimating genome size variation within human and Arabidopsis using k-mer frequencies. Bioinformatics 34:550-557.

Vermeij GJ. 1991. Anatomy of an invasion: the trans-Arctic interchange. Paleobiology 17:281-307.

Vurture GW, Sedlazeck FJ, Nattestad M, Underwood CJ, Fang H, Gurtowski J, Schatz MC. 2017. GenomeScope: fast reference-free genome profiling from short reads. Bioinformatics 33:2202-2204.

Wares JP, Cunningham CW. 2001. Phylogeography and Historical Ecology of the North Atlantic Intertidal. Evolution 55:2455-2469.
